# Interactomic analysis reveals a new homeostatic role for the HIV restriction factor TRIM5α in mitophagy

**DOI:** 10.1101/2021.08.20.457143

**Authors:** Bhaskar Saha, Michelle Salemi, Geneva L Williams, Michael L Paffett, Brett Phinney, Michael A Mandell

## Abstract

The protein TRIM5α has multiple roles in anti-retroviral defense, but the mechanisms underlying TRIM5α action are unclear. Here, we used an APEX2-based proteomics approach to identify TRIM5α-interacting proteins. Analysis of the TRIM5α interactome found proteins participating in a wide variety of cellular functions including regulating antiviral signaling pathways. We used this data set to uncover a novel role for TRIM5α in mitophagy, an autophagy-based mode of mitochondrial quality control that is compromised in multiple human diseases. Mitochondrial damage triggered the relocalization of TRIM5α to ER-mitochondria contact sites where TRIM5α colocalized with markers of autophagy initiation and autophagosome biogenesis. Furthermore, we found that TRIM5α knockout attenuated both Parkin-dependent and Parkin-independent mitophagy by preventing the recruitment of autophagy regulators FIP200 and ATG13 to unhealthy mitochondria. Finally, TRIM5α knockout cells showed reduced mitochondrial function under basal conditions and were more susceptible to uncontrolled immune activation and cell death in response to mitochondrial damage than were wild type cells. Taken together, our studies have identified a homeostatic role for a protein previously recognized exclusively for its antiviral actions.

## INTRODUCTION

The protein TRIM5α (tripartite motif containing protein 5 isoform α; TRIM5) has evolved as a cell-intrinsic antiviral factor capable of restricting diverse RNA viruses (Chiramel, Meyerson et al., 2019, Ganser-Pornillos & Pornillos, 2019) and endogenous retroelements (Volkmann, Wittmann et al., 2020). The best-studied antiviral activity of TRIM5 is its capacity to potently block retroviral infection at a stage after the entry of the virus but prior to integration of the proviral DNA into the host cell genome (Ganser-Pornillos & Pornillos, 2019). This mode of retroviral restriction correlates with the ability of TRIM5 to recognize the assembled retroviral capsid (Stremlau, Perron et al., 2006). While TRIM5 from rhesus macaques (RhTRIM5) efficiently recognizes the HIV-1 capsid in a manner that can block infection, human TRIM5’s (HuTRIM5) ability to recognize HIV-1 is substantially reduced and consequently HuTRIM5 has been considered ineffectual against HIV-1. Nevertheless, epidemiological studies have demonstrated that different alleles of TRIM5 are associated with altered risk of HIV-1 infection and disease progression in people (Cloherty, Rader et al., 2021). Since HuTRIM5 cannot carry out capsid-specific restriction of HIV-1, the human genetics studies imply that additional TRIM5-dependent mechanisms may be important for determining HIV-1 outcomes in humans.

In fact, recent studies have identified three novel TRIM5 actions whereby HuTRIM5 can impact the outcome of HIV-1 infection *in vitro*. First, activation of human cells with type I interferon enables HuTRIM5 to restrict HIV-1 (Jimenez-Guardeno, Apolonia et al., 2019, OhAinle, Helms et al., 2018). The enhanced potency of HuTRIM5 is linked to increased interactions between TRIM5 and the immunoproteasome which consists of different protein subunits than conventional proteasomes (Jimenez-Guardeno et al., 2019). Second, TRIM5 can activate antiviral signaling pathways leading to the production of cytokines including type I interferon and the generation of an antiviral state (Fletcher, Vaysburd et al., 2018, Merindol, El-Far et al., 2018, Pertel, Hausmann et al., 2011, Saha, Chisholm et al., 2020). The signaling function of TRIM5 at least partially depends on TRIM5’s interaction with the TGF-β activated kinase 1 (TAK1) protein complex (Pertel et al., 2011). Finally, TRIM5 interacts with multiple proteins having key roles in autophagy (Keown, Black et al., 2018, Mandell, Jain et al., 2014, O’Connor, Pertel et al., 2010, Ribeiro, Sarrami-Forooshani et al., 2016). Autophagy is a homeostatic mechanism of cellular waste management in which cytoplasmic contents (e.g. damaged mitochondria) are sequestered in a vesicle termed an autophagosome that ultimately fuses with lysosomes for elimination (Deretic, 2021, Youle, 2019). TRIM5’s connections to the autophagy pathway underlie the relative resistance to HIV-1 infection seen in human Langerhans cells(Ribeiro et al., 2016). Since autophagy has a plethora of functions in controlling cell survival, cellular metabolism, and innate immune or inflammatory responses (Deretic, 2021), it is likely that TRIM5 could contribute to these functions in a way that impacts the outcome of viral infection.

Clearly, an understanding of TRIM5’s protein-protein interactions has been crucial to understanding its numerous functions, yet questions about TRIM5 mechanisms remain elusive; introducing the possibility that novel TRIM5 activities exist. The premise of this study is that identifying TRIM5 interacting proteins (the TRIM5 interactome), will provide fundamental knowledge required for addressing these questions. To this end, we used an APEX2-based proximity ligation system to study TRIM5 protein-protein interactions since this approach allows for the direct labeling of TRIM5-promiximal proteins in intact, living cells leading to a high degree of coverage and of labeling specificity (Rhee, Zou et al., 2013). Using this approach, we identified more than 300 proteins with diverse functions as putative TRIM5 interactors. We noted an unexpected enrichment of mitochondrial proteins in the TRIM5 interactome which led us to identify a novel homeostatic role for TRIM5 in the autophagy-based elimination of dysfunctional mitochondria (mitophagy). TRIM5 acted as an assembly scaffold linking markers of damaged mitochondria with upstream autophagy regulators at the site where autophagosome assembly initiates. TRIM5-dependent mitophagy was crucial for preventing inflammation and cell death triggered by mitochondrial damage. In addition to identifying a key role for TRIM5 in mitophagy, our definition of the TRIM5 interactome will enable future discoveries of TRIM5 functions and mechanisms in and beyond TRIM5 actions in antiviral defense.

## RESULTS

### Proteomic definition of the TRIM5 interactome

We chose to use the APEX2 system to perform an unbiased identification of TRIM5 interacting partners (Trinkle-Mulcahy, 2019). APEX2 is an engineered form of soybean ascorbate peroxidase (Lam, Martell et al., 2015). In the presence of hydrogen peroxide, APEX2 catalyzes the conversion of membrane-permeable biotin-phenol to membrane-impermeant biotin-phenoxyl radical. This compound is highly reactive, and in cells it almost immediately reacts with electron-rich amino acid residues within a radius of less than 20 nm leading to proteins in the very near proximity to APEX2 or to APEX2-fusion proteins being covalently labeled with biotin (Rhee et al., 2013). Biotinylated proteins or protein complexes can then be isolated from cell lysates with streptavidin-coated beads and identified by LC-MS/MS (Jia, Abudu et al., 2018, Lam et al., 2015, Le Guerroue, Eck et al., 2017, Rhee et al., 2013) (Fig. 1A). We designed our screen to allow us to detect TRIM5-proximal proteins under homeostatic conditions and to detect potential changes in the TRIM5-interactome stimulated by TRIM5 binding to assembled retroviral cores that are released into the cytoplasm following fusion of the viral envelope with host cell membranes shortly after infection (Fig. 1B). For these experiments, we chose to use the well-established model of HIV-1 restriction by rhesus macaque TRIM5 since rhesus, but not human, TRIM5 is capable of binding to the HIV-1 viral cores (Ganser-Pornillos & Pornillos, 2019).

**Figure 1.**
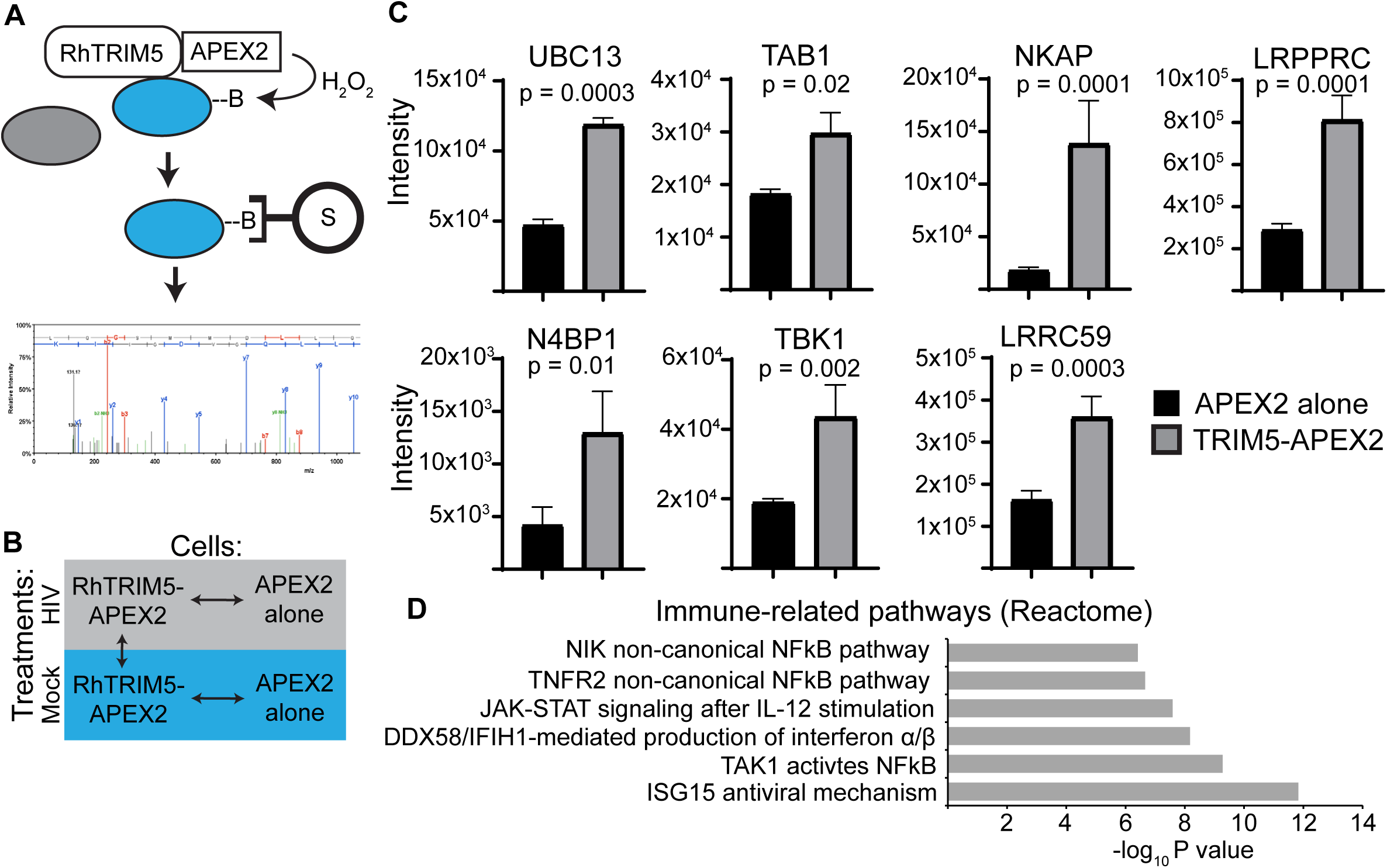
Proximity biotinylation screen for TRIM5 proximal proteins. **(A)** Schematic of APEX2-based labeling, purification, and identification of proteins in the very near vicinity (within 20 nm) of TRIM5. HEK293T cells stably expressing rhesus (Rh) TRIM5-APEX2-V5 or APEX2-V5 alone were cultured in the presence of biotin-phenol and then pulsed with H2O2 for 1 minute to induce covalent biotinylation of nearby proteins (within 20 nm). Biotinylated proteins were then purified on streptavidin-coated beads and subjected to mass spectrometry-based analysis. B, biotin; S, streptavidin. **(B)** Diagram of the comparisons performed for analysis of mass spectrometry data. Biotinylation reactions and cell lysis was performed 3 hours after synchronized infection of cells with VSV-G pseudotyped HIV-1 (HIV) or mock infection. Three independent biological replicates were analyzed per treatment. Data for the comparisons indicated by the arrows are found in Table S1-S3. **(C)** Intensity measurements of selected immune-related proteins identified by mass spectrometry. Data: mean + SEM; *P* values determined by t test. **(D)** Gene set enrichment analysis of immune-related Reactome pathways enriched in TRIM5-APEX2 datasets relative to those from APEX2 alone.

We generated HEK293T cells stably expressing either human (Hu) or rhesus (Rh) TRIM5 fused at its C terminus to V5-tagged APEX2 (TRIM5-APEX2). As a negative control for our proteomics experiments, we also generated HEK293T cells stably expressing APEX2-V5 (APEX2; Fig. S1A). These cell lines displayed the expected susceptibility to infection with VSV G-pseudotyped HIV-1 in which the RhTRIM5-APEX2 robustly restricted HIV-1 infection relative to both the HuTRIM5-APEX2 and APEX2 alone cell lines (Fig. S1B, C). We also confirmed that over-expression of TRIM5-APEX2 fusions could activate NF-κB and AP1 transcription factors (Fig. S1D,E) in accordance with published reports (Fletcher et al., 2018, Pertel et al., 2011). The findings from these control experiments demonstrate that the TRIM5-APEX2 fusion proteins retained important biological activities of untagged TRIM5.

The results of our data-independent acquisition mass spectroscopic analysis are found in Table S1-S3. Principal component analysis demonstrated that the TRIM5-APEX2 samples (both HIV-infected and mock) grouped together and differed from the APEX2 alone samples (Fig. S1F). We identified 356 proteins that were more than 1.6-fold enriched in the TRIM5-APEX2 samples (*P* < 0.05; Table S1-S3). Encouragingly, among these proteins we identified two previously known TRIM5 interactors that are involved in TRIM5-dependent antiviral signaling: TGF-beta activated kinase binding protein 1/TAB1; and Ubiquitin conjugating enzyme E2 N/UBC13 (Fig. 1C). Consistent with TRIM5’s actions in innate immune signaling, we identified multiple proteins that have roles in regulating signaling pathways that terminate in activation of NF-κB or IRF-3 (Fig. 1C, Table S1). In fact, gene set enrichment analysis of the proteins enriched in the TRIM5 interactome identified several immune-related pathways to be over-represented (Fig. 1D). These results were consistent regardless of the presence of HIV-1 (Table S1-S3), and we were unable to detect robust differences between mock infected and HIV infected samples in these experiments (Fig. S1G). Coimmunoprecipitation analysis of selected putative TRIM5 interactors identified in our screen show that a high percentage of those tested were in protein complexes with HA-tagged TRIM5 in HeLa cells under homeostatic conditions (Fig. S1H; Fig. 2E-G). Of 13 ‘hits’ tested, 9 showed positive interactions with TRIM5-HA in these experiments (Fig. S1I). Together, these data help validate the findings from our APEX2-based identification of TRIM5 interacting proteins.

**Figure 2.**
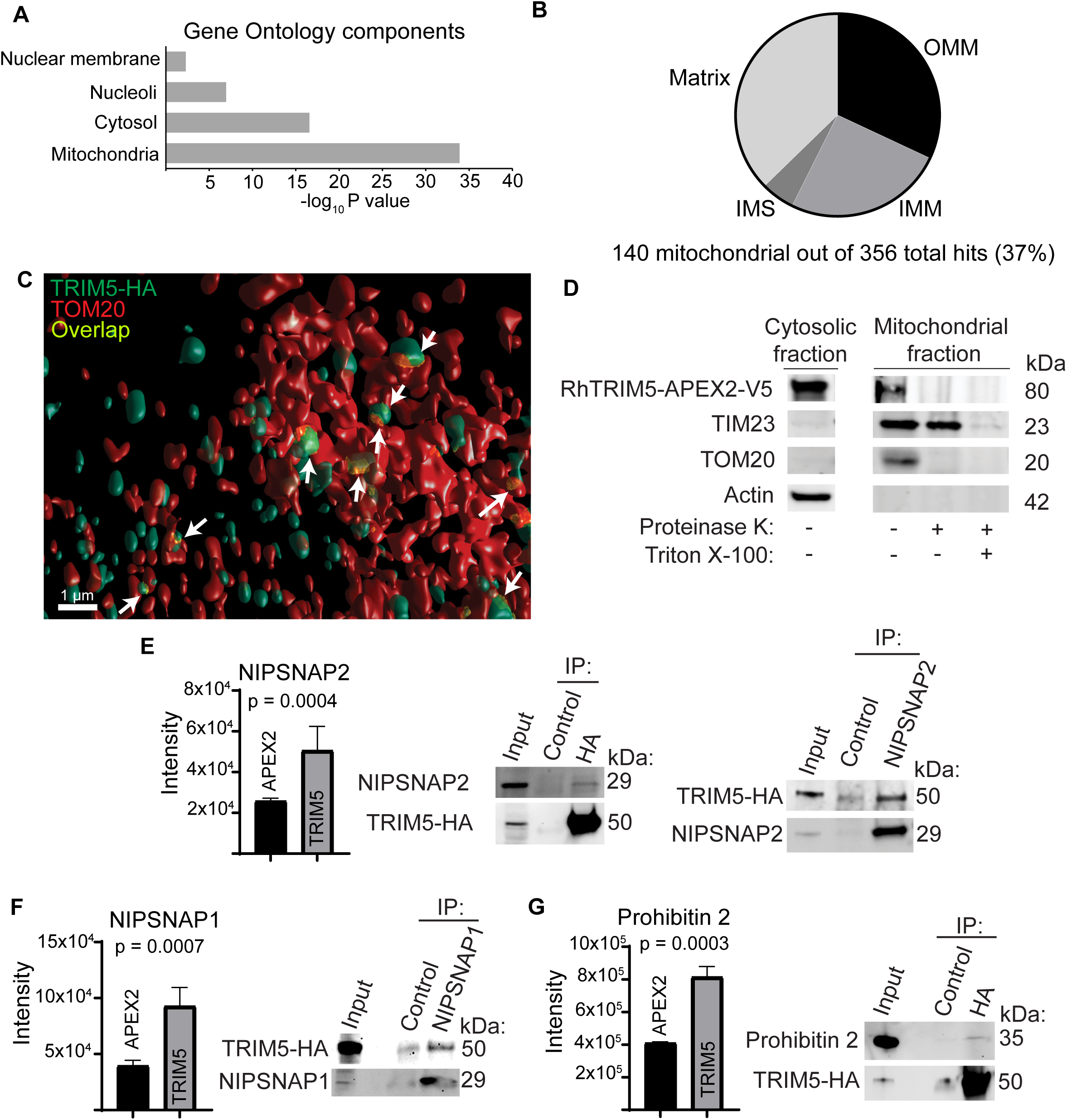
Mitochondrial localization of TRIM5 under basal conditions. **(A)** Cellular localization of TRIM5-proximal proteins based on gene ontology analysis. **(B)** Analysis of the sub-organellar localization of mitochondrial TRIM5 interactors based on gene ontology analysis. OMM, outer mitochondrial membrane; IMS, inner mitochondrial space; IMM, inner mitochondrial membrane. **(C)** 3D reconstruction of a deconvolved confocal micrograph showing a HeLa cell stained to detect TRIM5-HA (green) and the mitochondrial marker TOM20 (red). Overlapping volumes (yellow) are indicated by arrows. **(D)** Protease protection analysis of the mitochondrial localization of TRIM5. Mitochondria were purified from HEK293T cells stably expressing rhesus TRIM5-APEX2-V5 and treated or not with proteinase K in the presence or absence of detergent (Triton-X) prior to immunoblotting with the indicated antibodies. **(E-G)** Mass spectrometry- and coimmunoprecipitation-based analysis of interactions between TRIM5 and mitochondrial proteins that have been identified as mitophagy receptors. Graphs, intensity measurements of selected immune-related proteins identified by mass spectrometry. Data: mean + SEM; *P* values determined by t test. For coimmunoprecipitation experiments, lysates from HeLa cells stably expressing TRIM5-HA were immunoprecipitated as indicated prior to immunoblotting.

### The TRIM5 interactome is enriched in mitochondrial proteins

Previous studies have shown that TRIM5 primarily localizes to puncta in the cytoplasm (Campbell, Dodding et al., 2007) and can also be found in the nucleus (Diaz-Griffero, Gallo et al., 2011). However, gene ontology analysis of our mass spectroscopy data revealed an unexpected enrichment of mitochondrial proteins within the TRIM5 interactome (Fig. 2A). Further analysis of the mitochondrial proteins enriched in the TRIM5 interactome showed that TRIM5 was proximal to proteins designated as localizing to the outer mitochondrial membrane, the intermembrane space, the inner mitochondrial membrane and the mitochondrial matrix in previous APEX-based proteomics studies (Fig. 2B) (Hung, Lam et al., 2017, Hung, Zou et al., 2014, Rhee et al., 2013). We next used imaging and biochemical approaches to independently test the association between TRIM5 and mitochondria. In confocal microscopy experiments, cytoplasmic bodies of HA-tagged TRIM5 were often found to be very close to and likely in physical contact with structures positive for the outer mitochondrial membrane marker TOM20 (Fig. 2C; S2A). Remarkably, published correlative light electron cryotomography studies of TRIM5 localization also found TRIM5 structures that appeared to contact mitochondria (Carter, Mamede et al., 2020) in agreement with our data. Furthermore, TRIM5 was present in preparations of purified mitochondria (Fig. 2D). Protease treatment of intact purified mitochondria eliminated mitochondria-associated TRIM5 and TOM20 but not the inner mitochondrial membrane protein TIM23, indicating that TRIM5 was primarily associated with the cytoplasmic face of the outer mitochondrial membrane. Overall, these studies indicate a previously unappreciated association between TRIM5 and mitochondria. Given the key roles of mitochondria as sources of endogenous inflammation and as key orchestrators of innate antiviral signaling, we pursued further studies into what role(s) mitochondria-associated TRIM5 may play in the cell.

### TRIM5 interacts with mitochondrial proteins associated with mitophagy and relocalizes to mitochondria-ER contact sites in response to mitochondrial damage

Coimmunoprecipitation experiments confirmed that TRIM5 was in protein complexes with mitochondrial proteins NIPSNAP1, NIPSNAP2, Prohibitin 2, and LRPPRC in HeLa cell lysates (Fig. 2E-G; S1H,I). We also confirmed TRIM5-NIPSNAP2 interactions in HeLa cells by *in situ* proximity ligation assay (PLA), which indicates the presence of two proteins within 16 nm of each other by the formation of fluorescent puncta (Fig. S2B). Interestingly, NIPSNAP1/2, Prohibitin 2 (Fig. 2E-G) and SAMM50 (Fig. S2C) are reported to have roles in an autophagy-dependent mode of mitochondrial quality control referred to as ‘mitophagy’ (Abudu, Shrestha et al., 2021, Princely Abudu, Pankiv et al., 2019, Wei, Chiang et al., 2017). Mitophagy involves the selective removal of damaged or unwanted mitochondria, and aberrant mitophagy has been connected to multiple aging-related inflammatory conditions and cancers (Chen, Kroemer et al., 2020). In healthy mitochondria, the NIPSNAPs, Prohibitin 2, and SAMM50 are all primarily localized on the inside of the mitochondrion and away from the cytoplasm. However, mitochondrial damage increases their exposure to the cytoplasm where they are proposed to facilitate mitophagy by interacting with autophagosome membrane-associated proteins (LC3s and GABARAPs; the mammalian orthologues of yeast Atg8, mAtg8s) or the autophagy adaptor p62 (Abudu et al., 2021, Princely Abudu et al., 2019, Wei et al., 2017). Given TRIM5’s previously reported interactions with mAtg8s, p62, and additional components of the autophagy machinery (Keown et al., 2018, Mandell et al., 2014, Ribeiro et al., 2016), we investigated whether TRIM5 might be involved in mitophagy.

NIPSNAPs and Prohibitin 2 are both reported to contribute to a pathway referred to as “Parkin-dependent mitophagy” (Princely Abudu et al., 2019, Wei et al., 2017). In this pathway, the ubiquitin ligase Parkin relocalizes from the cytoplasm to the surface of damaged mitochondria. Once there, Parkin catalyzes the ubiquitylation of many outer mitochondrial membrane proteins and the subsequent recruitment of ubiquitin-binding autophagy adaptors including NDP52 and optineurin (Harper, Ordureau et al., 2018). A classic model of Parkin-dependent mitophagy involves inducing mitochondrial depolarization with carbonyl cyanide 3-chlorophenylhydrazone (CCCP) in HeLa cells expressing exogenous Parkin (HeLa cells lack Parkin expression). We used this model to determine if CCCP treatment could impact the mitochondrial localization of TRIM5 as a first test of whether TRIM5 contributes to mitophagy. As described previously, in untreated cells mCherry-tagged Parkin has a diffuse cytoplasmic localization and TRIM5 primarily localizes to small cytoplasmic puncta (Fig. 3A). However, two hours after CCCP treatment, Parkin has almost completely relocalized to numerous ovoid structures in the cell (presumably mitochondria; Fig. 3B and Fig. S3A). TRIM5’s localization also changes dramatically in response to CCCP treatment. Instead of localizing to small, round bodies as in untreated cells, in CCCP-treated cells TRIM5 forms elongated filaments. These filaments, which are often curved and sometimes have a branching morphology, appeared to intersect with and sometimes partially surround the Parkin-positive structures (Fig. 3B; S3A). Examination of these filaments in triple-labeling confocal microscopy experiments demonstrated that they closely associate with mitochondria (Fig. 3C; S3B). Interestingly, while many mitochondria showed labeling with both Parkin and TRIM5, the mitochondrial TRIM5 does not always localize to the same site as Parkin (Fig. 3C) and TRIM5-positive, Parkin-negative mitochondria are readily observed (Fig. S3B). These data indicate that TRIM5 and Parkin are independently recruited to damaged mitochondria and suggest that they may play distinct roles in mitophagy.

**Figure 3.**
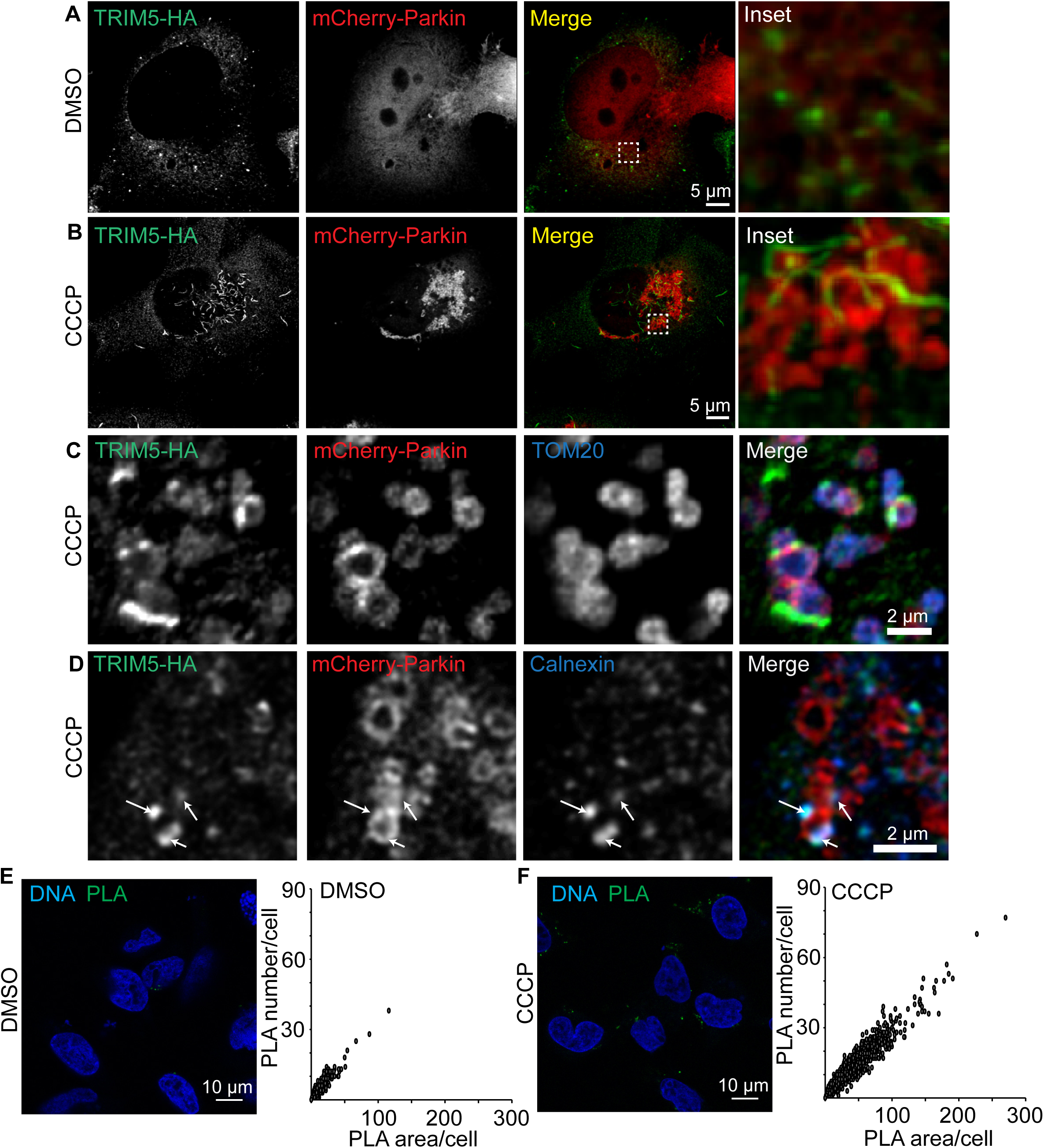
TRIM5 and autophagy proteins localize to ER-mitochondria contact sites in response to mitochondrial depolarization. HeLa cells stably expressing TRIM5-HA were transiently transfected with mCherry-Parkin prior to three-hour treatment with CCCP (20 µM) or vehicle control (DMSO). Cells were fixed, stained as indicated, and deconvolved confocal micrographs were acquired. **(A, B)** Localization of TRIM5 and mCherry-Parkin under control conditions (DMSO) and following mitochondrial uncoupling (CCCP). Dashed line shows the location of the zoomed-in region shown to the right. **(C)** Confocal analysis of the TRIM5-HA, mCherry-Parkin, and TOM20 following mitochondrial depolarization. **(D)** Representative images of cells treated as above and stained to detect TRIM5-HA (green), mCherry-Parkin (red) and ER marker Calnexin (D) in blue. Arrows indicate foci of colocalized green and blue signal on the outside of Parkin-decorated mitochondria. **(E,F)** Proximity ligation assay (PLA) probing the proximity of TRIM5-HA and the ER marker Calnexin in untreated cells (A) or in CCCP-treated cells (B). Micrographs show representative confocal images. Plots show data from high content imaging in which both the number of PLA puncta and the total cross-sectional area of PLA puncta were measured per cell. N > 5000 cells per treatment.

Based on their morphology, we hypothesized that the TRIM5 filaments seen in CCCP-treated cells localized to segments of the endoplasmic reticulum (ER). Indeed, we found substantial colocalization between TRIM5 and the ER marker Calnexin in CCCP-treated cells (Fig. 3D). Accordingly, analysis of our mass spectrometry results identified several ER-localized proteins and 13 ER-mitochondria contact site resident proteins (Hung et al., 2017) as being enriched in the TRIM5 interactome (Table S4). One of the ER-mitochondria contact site proteins identified in the TRIM5 interactome was Calnexin. *In situ* PLA confirmed that TRIM5 and Calnexin were in close proximity in untreated cells, and CCCP treatment substantially increased PLA signal (Fig. 3E) indicating that mitochondrial uncoupling recruits TRIM5 to ER. Taken together, our findings show that TRIM5 is recruited to ER-mitochondrial contact sites following mitochondrial damage.

### TRIM5 is required for Parkin dependent mitophagy in a cell type dependent manner

Previous studies have indicated that mitophagosome biogenesis originates from ER-mitochondria contact sites in both yeast and mammalian cells (Bockler & Westermann, 2014, Gelmetti, De Rosa et al., 2017, Yang, Li et al., 2020, Zachari, Gudmundsson et al., 2019). One of the earliest steps in autophagosome/mitophagosome formation is the generation of phosphatidyl inositol 3-phosphate (PI3P). DFCP1 is a PI3P binding protein that marks autophagy initiation sites. Following CCCP treatment, TRIM5 and GFP-DFCP1 colocalized with each other on the outside of Parkin-decorated mitochondria (Fig. 4A). Mitochondria-associated TRIM5 also colocalizes with endogenous FIP200 and ATG13, both key early players in autophagy regulation, in CCCP-treated cells (Fig. 4B, C). Typically, TRIM5 and autophagy factors localized to discrete points on the mitochondria rather than completely surrounding them (Fig. S4A,B). Importantly, colocalized ATG13 and TRIM5 was observed associated with mitochondria in CCCP-treated HeLa cells even in the absence of Parkin expression (Fig. S4C), suggesting that the recruitment of TRIM5 and autophagy regulatory machinery to damaged mitochondria is Parkin-independent.

**Figure 4.**
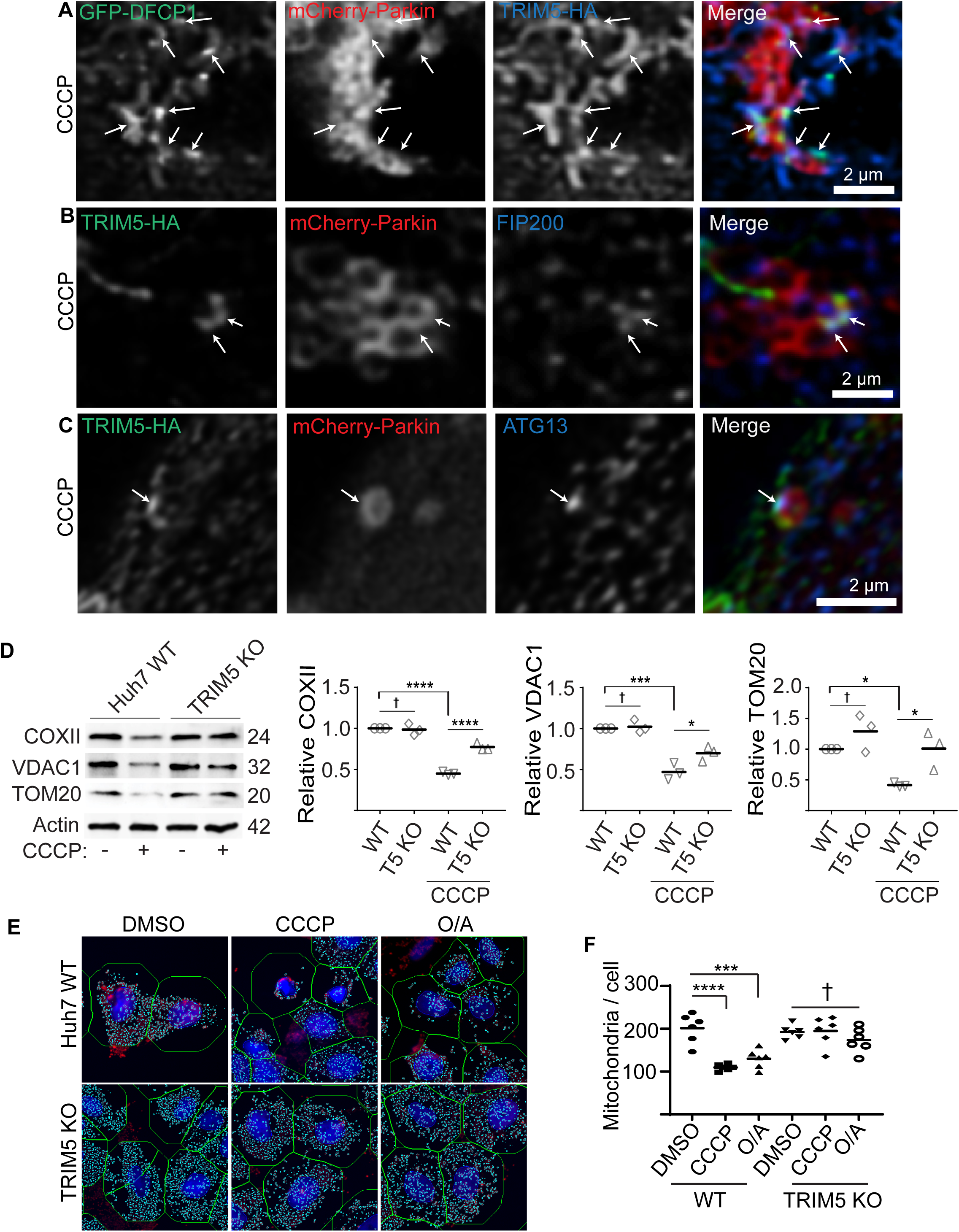
TRIM5 is required for mitophagy in response to mitochondrial depolarization. **(A)** HeLa cells stably expressing TRIM5-HA (blue) were transiently transfected with both mCherry-Parkin (red) and eGFP-DFCP1 (green) prior to 2 h CCCP treatment and confocal microscopy. Arrows indicate regions of TRIM5/DFCP1 colocalization that are associated with Parkin-labeled mitochondria. **(B, C)** Representative confocal images of cells treated as above and stained to detect TRIM5-HA (green), mCherry-Parkin (red) and the autophagy factors FIP200 (B) or ATG13 (C) in blue. Arrows indicate foci of colocalized green and blue signal on the outside of Parkin-decorated mitochondria. **(D)** Immunoblot-based analysis of mitochondrial protein abundance in lysates from WT and TRIM5 knockout Huh7 cells under basal conditions and following 24 h treatment with CCCP. The abundance of these proteins from three independent experiments are plotted relative to actin. Data: *, *P* < 0.05; ***, *P* < 0.001; ****, *P* < 0.0001, †, not significant by ANOVA. **(E, F)** High content imaging-based analysis of the abundance of mitochondrial nucleoids in WT and TRIM5 KO Huh7 cells treated with CCCP (10 µM) or oligomycin and antimycin (O/A; 10 µM and 5 µM, respectively) for 24 h prior to fixation and staining with an antibody that preferentially recognized mitochondrial DNA (red) and nuclear staining (blue). Left, representative images showing automatically defined nuclear and cell boundaries (blue and green rings, respectively) and mitochondrial nucleoids (aqua mask). The number of mitochondria per cell was determined for 2000 cells per experiment. Data: each point represents an independent experiment. Data: ***, *P* < 0.001; ****, *P* < 0.0001; †, not significant by ANOVA.

Since TRIM5 localized to apparent mitophagosome formation sites, we next tested its role in mitophagy using the Parkin-expressing HeLa cell model (Fig. S5). As expected, treatment of HeLa cells stably over-expressing YFP-Parkin with CCCP or the mitochondrial respiration inhibitors oligomycin/antimycin (O/A) induced mitophagy as measured by increased degradation of the mitochondrial proteins COXII and VDAC1. However, TRIM5 knockout partially reversed this effect (Fig. S5A, B). Similar results were seen when the abundance of the lipidated, autophagy-specific form of LC3 (LC3-II) were measured (Fig. S5C, D), indicating that TRIM5 contributes to, but is not required for, mitophagy in this model system. High content imaging experiments showed that TRIM5 knockout did not affect the kinetics of CCCP-dependent relocalization of Parkin to mitochondria (Fig. S5E, F) or CCCP-induced formation of ubiquitin puncta (Fig. S5G,H), suggesting that TRIM5 contributes to mitophagy either by acting downstream of Parkin-mediated ubiquitination or by participating in a complementary pathway.

If the latter of these two models is correct, then the relative importance of TRIM5-dependent mitophagy might be higher in an experimental system in which the expression of TRIM5 more closely matched the expression of Parkin. Unlike HeLa cells, the hepatocellular adenocarcinoma cell line Huh7 expresses endogenous Parkin (Kim, Syed et al., 2013), and so we tested the role of TRIM5 in mitophagy using these cells. CCCP treatment of WT Huh7 cells reduced the levels of mitochondrial proteins COXII, VDAC1, and TOM20 by ∼50%. This effect was almost completely lost in TRIM5 knockout Huh7 cells (Fig. 4D). Expression of HuTRIM5-APEX2 restored the ability of CCCP to induce COXII degradation in TRIM5 knockout Huh7 cells (Fig. S5I). High content imaging revealed that CCCP or O/A treatment strongly reduced the number of mitochondrial nucleoids per cell in WT Huh7 cells but these treatments had no impact on TRIM5 knockout Huh7 cells (Fig. 4E, F). These results show that TRIM5 is required for mitophagy in a cell-type dependent manner.

### TRIM5 bridges damaged mitochondria with upstream autophagy regulators

We next sought to define the mechanism underlying TRIM5 action in CCCP-induced mitophagy. Confocal microscopy indicated that cells lacking TRIM5 have a defect in autophagosomal targeting of mitochondria. In CCCP-treated WT Huh7 we were readily able to detect individual mitochondria that were surrounded by LC3B indicating their presence within autophagosomes (Fig. 5A). In contrast, we could not find any convincing examples of LC3B-enveloped mitochondria in TRIM5 knockout cells, despite seeing numerous LC3B-positive punctate structures.

**Figure 5.**
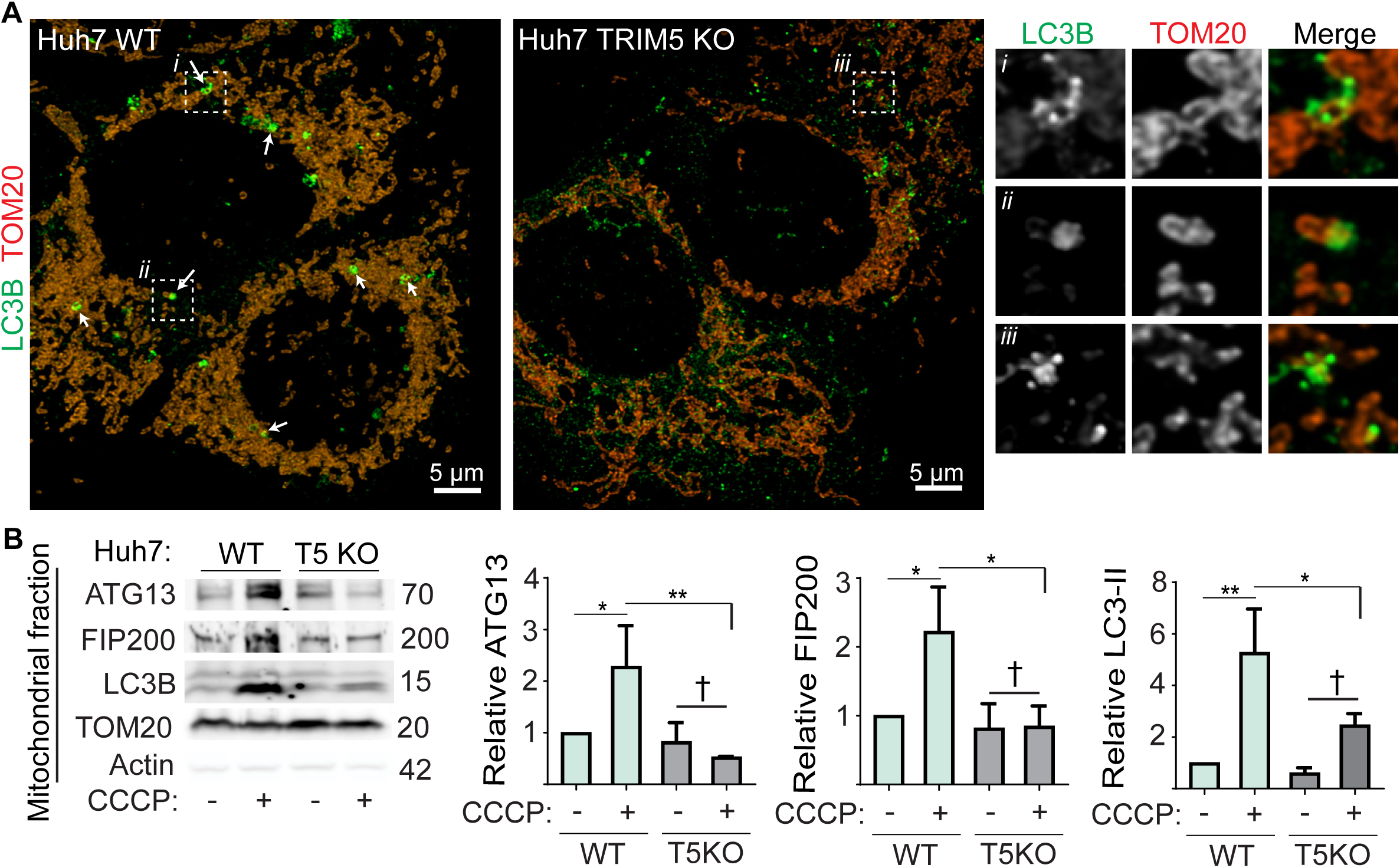
TRIM5 recruits upstream autophagy regulatory machinery to damaged mitochondria. **(A)** Confocal microscopic analysis of LC3B and TOM20 association with WT and TRIM5 KO Huh7 cells. Arrows indicate TOM20-positive structures (mitochondria) that are enveloped within LC3B-positive signaling. Zoomed-in images of the boxed regions are shown on the right. LC3B-surrounded mitochondria could not be identified in images from TRIM5 KO Huh7 cells. **(B)** Recruitment of autophagy machinery to mitochondria following CCCP treatment. Lysates from mitochondria purified from WT or TRIM5 Huh7 cells treated with CCCP (20 µM) or vehicle control (DMSO) for 6 hours were subjected to immunoblotting with the indicated antibodies. Plots show the abundance of the indicated proteins in the mitochondrial fractions relative to TOM20. Data: mean + SEM; *, *P* < 0.05; **, *P* < 0.01; †, not significant by ANOVA.

TRIM5, along with several other TRIM protein family members, has been shown to interact with multiple autophagy proteins including ULK1 (Chauhan, Kumar et al., 2016, Di Rienzo, Antonioli et al., 2019, Kimura, Jain et al., 2015, Mandell et al., 2014). ULK1, along with its binding partners ATG101, ATG13, and FIP200 are the most upstream components of the core autophagy machinery. Our data showed that TRIM5 colocalized with ATG13 and FIP200 on mitochondrial surfaces after CCCP treatment (Fig. 4B, C; S4A-C), suggesting that TRIM5 may act in mitophagy by recruiting these proteins to damaged mitochondria. We tested how TRIM5 knockout would impact the abundance of autophagy proteins that co-purify with mitochondria in CCCP-treated or untreated cells (Fig. 5B). In WT Huh7 cells, CCCP treatment increased the abundance of ATG13, FIP200, and the autophagosome-associated protein LC3B-II in mitochondrial fractions. Remarkably, we observed no enrichment of any of these proteins in mitochondrial fractions harvested from CCCP-treated TRIM5 knockout Huh7 cells. This result demonstrates that TRIM5 is required for the recruitment of upstream autophagy regulators to damaged mitochondria in cells expressing physiologically relevant levels of Parkin.

### TRIM5 acts independently of Parkin in mitophagy

While Parkin is required for the best studied mitophagy pathways, it has become clear that Parkin-independent mitophagic pathways exist and there are an increasing number of examples in which Parkin-independent mitophagy is an essential contributor to normal physiological function (Villa, Marchetti et al., 2018). We next tested whether TRIM5 might be important for mitophagic-responses to the iron chelator deferiprone (DFP) (Allen, Toth et al., 2013) or the anti-helminthic drug ivermectin (IVM) (Zachari et al., 2019) as both of these compounds are reported to induce Parkin-independent mitophagy. While DFP induced substantial degradation of mitochondrial proteins in HeLa cells, this effect was unchanged in TRIM5 knockout cells (Fig. S6A), suggesting that TRIM5 is dispensable for DFP-induced mitophagy. In contrast, we found that TRIM5 was required for IVM-induced degradation of COXII and VDAC1 in HeLa cells (Fig. 6A-D). We saw similar results when we performed this experiment in WT and TRIM5 knockout Huh7 cells, although in this case we also observed a role for TRIM5 in the IVM-induced degradation of TOM20 (Fig. S6B-E). Expression of TRIM5-APEX2, but not APEX2 alone, restored the ability of IVM to promote COX-II degradation in TRIM5 knockout HeLa cells (Fig S6F). Moreover, we found that TRIM5 knockout prevented the IVM-induced formation of LC3B-II in HeLa cells (Fig. 6E,F), suggesting that TRIM5 is required for autophagic responses to IVM.

**Figure 6.**
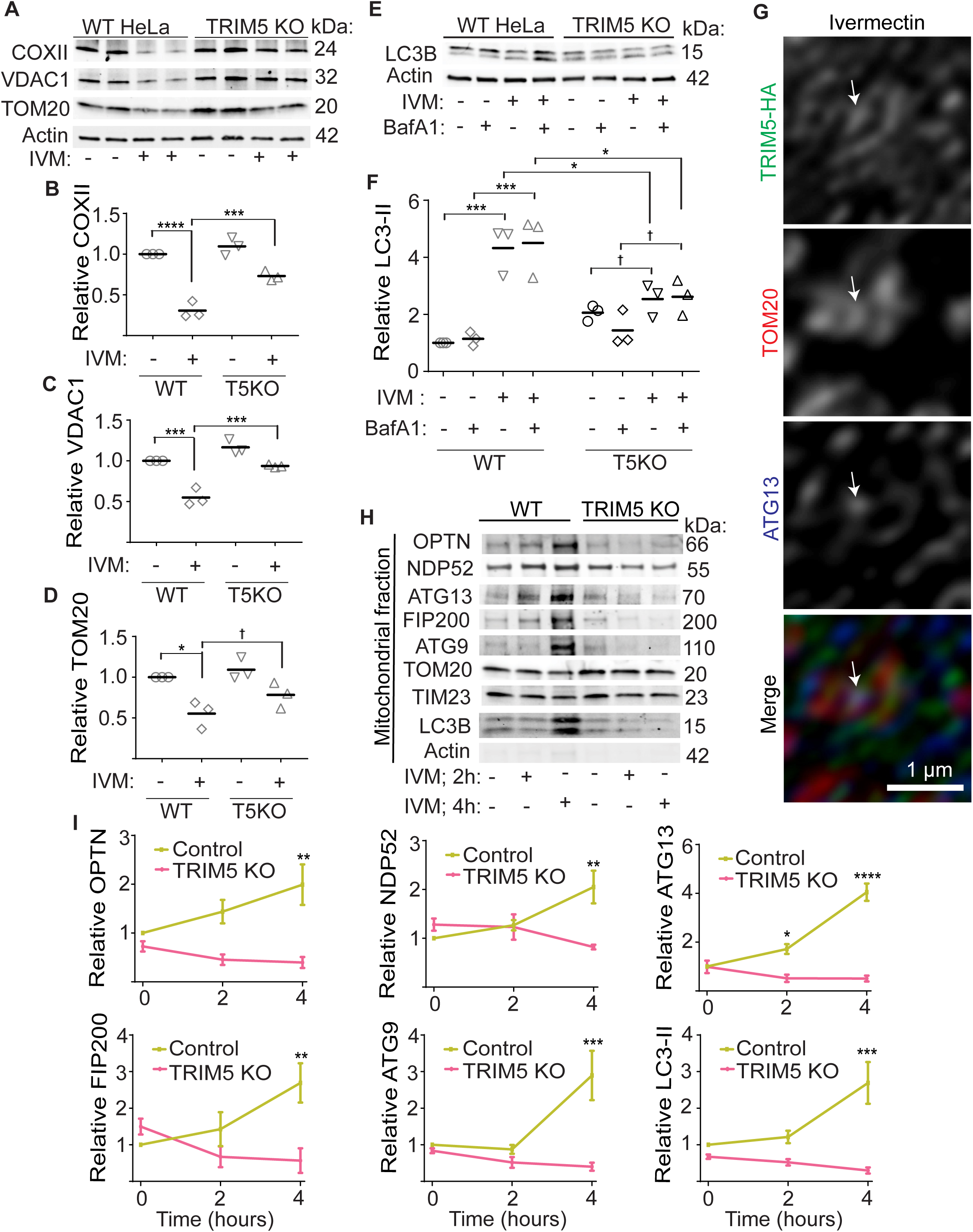
TRIM5 is required for Parkin-independent mitophagy stimulated by ivermectin. **(A-D)** Immunoblot analysis of the effect of TRIM5 knockout on ivermectin-stimulated degradation of COXII, VDAC1, and TOM20 in HeLa cells. Cells were treated or not with ivermectin (15 µM) for 24 hours prior to lysis and immunoblotting. The abundance of the proteins indicated in the graphs was determined relative to actin. Each data point represents an independent experiment. **(E, F)** Immunoblot analysis of the impact of TRIM5 knockout on the formation of lipidated LC3B (LC3-II) in response to ivermectin treatment (15 µM, 3 h) in the presence and absence of bafilomycin A1 (BafA1). Each data point represents an independent experiment. **(G)** Localization analysis of TRIM5-HA, ATG13, and TOM20 in HeLa cells following 2 h ivermectin treatment. Arrow indicates a site on a mitochondrion to which both TRIM5 and ATG13 localize. **(H, I)** The impact of TRIM5 knockout on the ivermectin-induced recruitment of the indicated autophagy-related proteins to mitochondria. Intact mitochondria were isolated from WT or TRIM5 knockout HeLa cells that treated with ivermectin for 2 or 4 hours or with vehicle control (DMSO) for 4 hours. Plots shown mean ± SEM. N = 3 experiments. *, *P* < 0.05; **, *P* < 0.01; ***, *P* < 0.001; ****, *P* < 0.0001; †, not significant by ANOVA.

Our data indicate that TRIM5 plays a similar role in IVM-induced mitophagy as what we described above for mitophagy induced by the mitochondrial uncoupling agent CCCP. Confocal imaging experiments revealed that TRIM5 localized to structures reminiscent of ER that were closely associated with mitochondria in IVM-treated HeLa cells and that showed colocalization with ATG13 (Fig. 6G). Mitochondrial fractionation experiments revealed that TRIM5 was required for the recruitment of autophagy factors to mitochondria after IVM treatment. Whereas the proteins optineurin, NDP52, ATG13, FIP200, ATG9, and LC3B were strongly enriched in mitochondrial fractions by 4 hours after IVM treatment, we observed no recruitment of these proteins to mitochondria in TRIM5 knockout cells (Fig. 6H, I). A previous study by Zachari et al showed that FIP200 was the earliest autophagy factor to be recruited to mitochondria in IVM treated cells and that FIP200 was required for the subsequent recruitment of other autophagy factors (Zachari et al., 2019). Since TRIM5 knockout prevented FIP200 recruitment, our data show that TRIM5 acts very early in mitophagy responses to IVM-induced mitochondrial damage. Taken together, these data show that TRIM5 plays an early role in some, but not all, Parkin-independent mitophagy mechanisms, likely by linking FIP200 complexes to damaged mitochondria.

### TRIM5-mediated mitophagy protects against mitochondrial damage and excessive inflammation

We next carried out metabolic profiling experiments to determine if TRIM5 was required for the maintenance of mitochondrial health. We found that WT Huh7 cells had a higher oxygen consumption rate (OCR) and had a higher capacity to carry out mitochondrial ATP synthesis than did TRIM5 knockout cells (Fig. 7A-D). In these experiments, we used IVM as a positive control for inducing mitochondrial damage (Zachari et al., 2019). We found that IVM treatment of WT Huh7 cells phenocopied what was seen in TRIM5 knockout Huh7 cells with or without IVM treatment. This result indicates that TRIM5 deficiency results in mitochondrial impairment even in the absence of mitophagy inducing stimuli and is consistent with our findings that TRIM5 plays an important role in mitophagy induced by mitochondrial damage. Furthermore, the fact that mitochondrial health was impaired in TRIM5 knockout cells without treatment hints that TRIM5 may also contribute to basal mitophagy.

**Figure 7.**
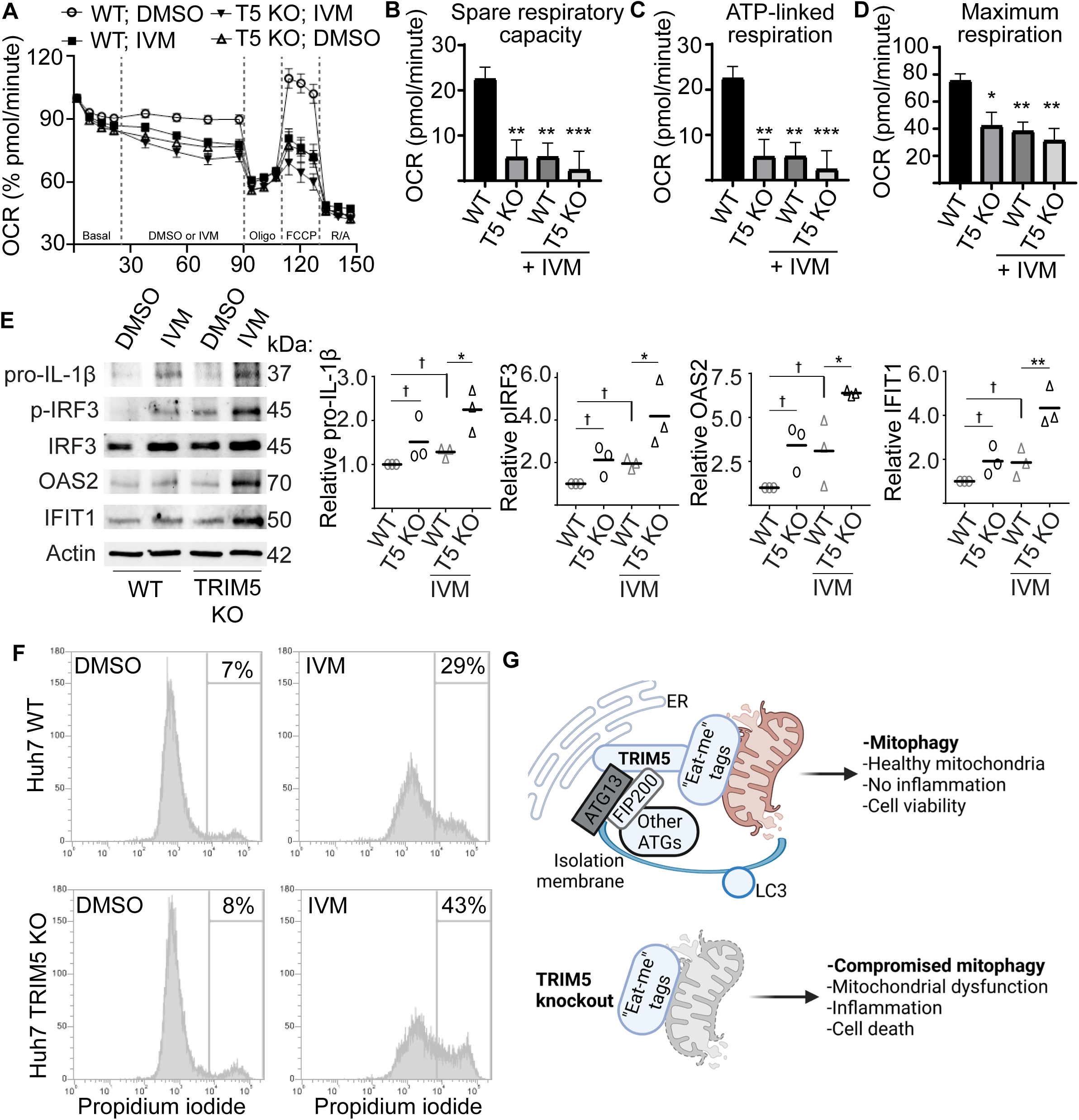
TRIM5 protects cells from excessive inflammation and cell death in response to mitochondrial damage. **(A-D)** Seahorse analysis of oxygen consumption rate (OCR) in WT and TRIM5 knockout Huh7 cells in response to ivermectin (IVM; 1.5 µM) or DMSO alone. **(E)** Immunoblot analysis of immune- and inflammation-related proteins in WT or TRIM5 knockout Huh7 cells following a 24 hour treatment with 15 µM IVM or DMSO control. Plots show the abundance of the indicated protein relative to a loading control (actin). Each data point represents an independent experiment. *, *P* < 0.05; †, not significant by ANOVA. **(F)** The impact of IVM treatment (20 µM; 24 hours) on the viability of WT and TRIM5 knockout Huh7 cells. The intensity of propidium iodide labeling, indicative of plasma membrane disruption, was determined in single cells by flow cytometry. **(G)** Schematic illustration of TRIM5 action in mitophagy. Top, TRIM5 interacts with mitophagy “eat-me” tags like NIPSNAP2 and Prohibitin 2 and also with ULK1 complex components ATG13 and FIP200 and other autophagy proteins (ATGs). TRIM5 recruits the autophagy machinery to damaged mitochondria at ER-mitochondria contact sites from which autophagy initiation emanates. TRIM5 contributes to mitophagic elimination of damaged mitochondria, resulting in increased mitochondrial and cellular health. Bottom, in the absence of TRIM5, damaged mitochondria are not removed by mitophagy. Consequently, TRIM5 knockout cells are more susceptible to inflammatory responses and cell death in response to mitochondrial damage.

If not removed from cells by mitophagy, damaged mitochondria can be important endogenous sources of immune activating DAMPs and PAMPs and can lead to cell death. We thus tested whether TRIM5’s actions in mitophagy are connected to its actions in regulating innate immunity and in antiviral defense. We found that 24 h treatment of WT Huh7 cells with IVM modestly increased the protein levels of active phosphorylated IRF3, the interferon-stimulated genes OAS2 and IFIT1, and the inactive form of the pro-inflammatory cytokine IL-1β (Fig. 7E). Importantly, the impact of IVM treatment on the expression of these immune-related proteins was substantially more pronounced in TRIM5 knockout Huh7 cells, suggesting that the defect in mitophagy in these cells is connected with enhanced inflammation. We also saw that TRIM5 knockout Huh7 cells were more susceptible to cell death following exposure to IVM than were WT Huh7 cells (Fig. 7F; Fig. S7A). Lastly, we tested whether TRIM5’s activities as a retroviral restriction factor were connected to its roles in mitophagy. While human TRIM5 is generally incapable of recognizing the capsid of HIV-1, a single amino acid substitution in the HIV capsid (P90A CA) renders the virus sensitive to HuTRIM5-based restriction (Kim, Dauphin et al., 2019, Selyutina, Persaud et al., 2020). Furthermore, the P90A CA virus can stimulate TRIM5-based antiviral signaling (Saha et al., 2020). However, infection of YFP-Parkin expressing HeLa cells with VSV-G pseudotyped HIV-1 P90A CA did not decrease the abundance of mitochondrial COXII, VDAC1, or TOM20 nor did it increase the abundance of autophagy marker LC3B-II (Fig. S7B). We also tested whether pre-treatment of cells with mitochondrial damaging agents CCCP or IVM could the impact ability of RhTRIM5 to potently restrict WT HIV-1 but we did not observe any effect (Fig. S7C). Taken together, our results show that TRIM5-dependent mitophagy is required to prevent excessive inflammation and cell death yet appears to be disconnected from TRIM5’s actions in retroviral restriction.

## DISCUSSION

In this study, we identified proteins proximal to or interacting with the HIV restriction factor TRIM5α and used these data to identify a novel role for TRIM5 in maintaining mitochondrial health through the autophagic removal of damaged mitochondria, a process termed ‘mitophagy’. In the absence of TRIM5, overall mitochondrial health is reduced and cellular sensitivity to some mitochondrial damaging agents such as ivermectin increase, resulting in increased inflammation and cell death. In contrast to previous studies that have focused on TRIM5’s specialized activities in protecting cells against viral infection, our study establishes that TRIM5 also has important homeostatic roles beyond antiviral defense that may be of relevance to the ever increasing number of pathological conditions associated with aberrant mitophagy.

The first indication that TRIM5 could be linked to mitochondria came from our finding that mitochondrial proteins were unexpectedly over-represented in the TRIM5 interactome (Table S1-S4). Among these mitochondrial TRIM5 interactors were multiple proteins that were previously identified as mitophagy ‘eat me’ tags including NIPSNAP1/2, Prohibitin 2, and SAMM50. Under conditions of mitochondrial stress, these proteins are presented on the cytoplasmic face of the outer mitochondrial membrane where they can flag damaged mitochondria for autophagic elimination through interactions with cytosolic autophagy proteins such as LC3/GABARAP, p62, and ALFY (Abudu et al., 2021, Princely Abudu et al., 2019, Wei et al., 2017). TRIM5 is known to interact with these and other autophagy proteins (Mandell et al., 2014), and our data show that TRIM5 is required for the recruitment of autophagy proteins to damaged mitochondria (Fig. 7G). Notably, we found that the recruitment of the autophagy proteins ATG13 and FIP200 to mitochondria following treatment with mitochondrial damaging agents was largely abrogated in TRIM5 knockout cells. FIP200 and ATG13 are components of the ULK1 complex which is required for the earliest steps in autophagosome or mitophagosome formation (Zachari & Ktistakis, 2020), suggesting that TRIM5 is an early upstream factor in mitophagy. Accordingly, TRIM5 knockout not only prevented autophagic degradation of mitochondrial components but it also prevented autophagosome formation in response to both IVM and CCCP.

Under normal conditions, TRIM5 localizes to cytoplasmic puncta, some of which intersect with mitochondria. Treatment of cells with mitochondrial damaging agents triggers a relocalization of TRIM5 to the ER, including regions of the ER that are in close contact with mitochondria. It is at these sites that TRIM5 colocalizes with ATG13 and FIP200 and also with DFCP1, a marker of autophagosome initiation sites. This makes sense, as ER-mitochondria contact sites have emerged as having key roles in autophagosome/mitophagosome biogenesis (Zachari & Ktistakis, 2020). Interestingly, several proteins known to reside at the ER or at ER-mitochondrial contact sites were found in the TRIM5 interactome (Table S1-S4). One of these, Atlastin 2, was recently shown to be required for the assembly of active ULK1 complexes to the ER and for the formation of autophagy initiation sites in response to amino acid starvation (Liu, Zhao et al., 2021). Whether TRIM5 and Atlastin 2 cooperate in recruiting or retaining ULK1 complexes to ER-mitochondrial contact sites or whether TRIM5 has other roles in assembling or activating the autophagy machinery at these sites requires further investigation.

While multiple distinct mechanisms for mitophagy exist in the literature (Zachari & Ktistakis, 2020), historically these mechanisms have been segregated based on their dependency on the ubiquitin ligase Parkin. Our data showed an involvement of TRIM5 in both Parkin-dependent mitophagy experimentally stimulated by treatment of cells with the compounds CCCP or oligomycin/antimycin A and in Parkin-independent mitophagy stimulated by ivermectin. In Parkin-dependent mitophagy, Parkin is recruited to damaged mitochondria from the cytoplasm by interacting with phosphorylated ubiquitin on the mitochondrial surface. Parkin then catalyzes the ubiquitylation of multiple outer mitochondrial membrane proteins which allows for the recruitment of ubiquitin-binding autophagy adaptors. Our data indicate that TRIM5 acts through a parallel and complementary pathway. Parkin was not required for TRIM5 localization to mitochondria, and we found TRIM5-ATG13 double-positive structures localized to mitochondria in Parkin-deficient HeLa cells. Likewise, TRIM5 knockout did not impair the ability of Parkin recruitment to and ubiquitylation of damaged mitochondria. Furthermore, it appears that a cell’s relative dependency on Parkin-dependent or TRIM5-dependent pathways can be variable, as TRIM5 knockout only modestly impacted mitophagy in a setting of Parkin over-expression but robustly impacted mitophagy in Huh7 cells that express endogenous Parkin. Thus, TRIM5-dependent mitophagy and Parkin-dependent mitophagy pathways can independently operate to rid cells of unwanted and potentially hazardous damaged mitochondria. Similarly to Parkin-dependent mitophagy, ubiquitylation of mitochondrial proteins is also an early step in Parkin-independent mitophagy stimulated by ivermectin (Zachari et al., 2019). An important question to be addressed by future investigation is whether the ubiquitin ligase activity of TRIM5 is required for TRIM5-dependent mitophagic pathways and, if so, what proteins are ubiquitylated by TRIM5 in response to mitochondrial damage. Importantly, our data show that TRIM5 is not required for all mitophagic pathways as TRIM5 knockout did not impair mitophagy driven by iron depletion resulting deferiprone (DFP) treatment. However, TRIM5 is a part of the large TRIM family of proteins consisting of ∼70 members in the human genome, and it is possible that other TRIM(s) contribute to mitophagic pathways induced by DFP or other stimuli.

In addition to uncovering a novel TRIM5-dependent mitophagy pathway, our identification of the TRIM5 interactome is likely to enable the elucidation of hitherto enigmatic TRIM5 mechanisms and may lead to the discovery of additional new roles for TRIM5. Although our data presented here shows that the loss of TRIM5 can enhance the expression of interferon stimulated genes in response to mitochondrial damage (presumably by disabling the cell’s ability to carry out the mitophagic removal of multiple mitochondria-associated DAMPs (Rai, Janardhan et al., 2021)), previous studies have shown that activation of TRIM5 by retroviral core engagement can trigger the expression of type I interferons (Fletcher et al., 2018, Merindol et al., 2018, Saha et al., 2020). While TRIM5 signaling is known to activate NF-κB and AP1 transcription factors (Pertel et al., 2011), the expression of type I interferon also requires activation of IRF-family transcription factors (e.g. IRF-3). Our proteomics data showed that several proteins known to regulate pathways upstream of IRF-3 (e.g. TBK1, LRPPRC, FAF2/UBXN3B, LRRC59 (Louis, Burns et al., 2018, Refolo, Ciccosanti et al., 2019, Xian, Yang et al., 2020, Yang, Wang et al., 2018) were enriched in the TRIM5 interactome (Table S1-S3), raising the possibility that these proteins are impacted by activated TRIM5 in a way that would trigger type I interferon expression. The role of these proteins in TRIM5-dependent immune activation and the establishment of an antiviral state will be a focus of future study.

In conclusion, our study has defined the interactome of TRIM5, an important antiviral restriction factor, and has uncovered a previously unanticipated homeostatic function of TRIM5 in preventing excessive inflammation and cell death in response to mitochondrial damage.

## Acknowledgements

This work was supported by P20GM121176 and R01AI155746 to M.A.M from the US National Institutes of Health. We thank Dr. Wade Johnson for technical expertise at the University of New Mexico Cancer Comprehensive Center Flow Cytometry Shared Resource which is partially supported by P30CA118100 form the NIH. We thank Dr. Ed Campbell (Loyola University Chicago) for sharing reagents and Dr. Jingyue Jia and Dr. Suresh Kumar for critically reading the manuscript. Biorender software was used for to generate graphics.

## Author contributions

B.S., M.S., G.L.W., and M.A.M performed research; M.L.P. shared technical expertise; B.S., M.S., B.P., and M.A.M designed research and analyzed data; B.S. and M.A.M. wrote the paper. The authors declare that they have no conflict of interest.

## MATERIALS AND METHODS

### Cell culture

HEK293T, HeLa, and Huh7 cells were obtained from the American Type Culture Collection (ATCC) and grown in Dulbecco’s modified Eagle’s medium (Life Technologies,11965126) supplemented with 10% fetal bovine serum (FBS, Life technologies, 26140-079), 100 U/ml penicillin and 100 µg/ml streptomycin at 37°C in a 5% CO2 atmosphere. HeLa cells stably expressing HA-tagged RhTRIM5α and HA-tagged HuTRIM5α were obtained from NIH AIDS reagents and were maintained in the above media supplemented with 1 µg/ml puromycin. YFP-PARKIN stably expressing HeLa were generated by Dr. Richard J. Youle (National Institutes of Health, USA). TRIM5 knockout (KO) Huh7, HeLa and YFP-PARKIN HeLa cells were generated by transduction with lentiCRISPRv2-based lentiviruses followed by 2-4 weeks of culturing in medium containing 200 µg/ml hygromycin. Knockout lines were confirmed by immunoblot and by confirmations of altered susceptibility to TRIM5-sensitve retroviruses. APEX2-V5, RhTRIM5-APEX2-V5 and HuTRIM5-APEX2-V5 stable overexpression in HEK293T, and in HeLa or Huh7 TRIM5 KO lines were achieved by viral transduction followed by 14-21 days of culturing in medium containing the selective antibiotic (1 µg/ml puromycin) before expression of the target gene was confirmed by western blotting.

### Generation of single-cycle lentivirus or retrovirus for stable cell line construction or infection assays

Viral particles for the generation of stable overexpressed cell lines were produced by co-transfection of pLEX_307 (a gift from David Root, Addgene plasmid # 41392) containing the target gene, psPAX2 and pMD2.G at the ratio of 1:1:1 in HEK293T cells using ProFection Mammalian Transfection System (Promega, E1200), medium was changed 16h post transfection and virus containing supernatant was harvested 48h later, clarified by centrifuging for 5 min at 1200 rpm, 0.45 µm-filtered (Millipore, SE1M003M00), diluted with full medium at 1:1 ratio and used to transduce target cells for 48 h.

Viral particles for the generation of knockout cell lines were produced by transfecting HEK293T cells with a lentiviral vector, lentiCRISPRv2 carrying both Cas9 enzyme and a guide RNA targeting specific gene together with the packaging plasmids psPAX2 and pMD2.G at the ratio of 10 µg, 10 µg and 10 µg/10 cm dish. We also used this approach to generate the HIV-1 CA P90A pseudoviruses. Lentiviral particles were harvested from supernatants as mentioned above and HEK293T, Huh7, and YFP-PARKIN HeLa cells were infected in the presence of polybrene for 48h in 6 cm dishes.

In this study, virus infection experiments were performed using VSV-G pseudotyped single-cycle HIV-1 produced by transfection of HEK293T cells. 24h before transfection, 2.2×10^6^ HEK293T cells were seeded in 10 cm dishes. Two-part single cycle VSV-G-pseudotyped HIV-1 (NL43 strain) was collected from the supernatants of HEK293T cells transfected with plasmids encoding VSV-G and HIV-1 lacking the Env gene (Env-, VPR-, Nef+, IRES-GFP) at 1:2 ratios. 16h post transfection, the culture media was changed, viral supernatant was harvested at 72h, clarified by centrifuging for 5 min at 1200 rpm, passed through a 0.45 µm filter and stored at - 80°C.

### Cloning and transfection

pDest40-APEX2-V5 and pLEX_307-APEX2-V5; pDest40-RhTRIM5-APEX2-V5 and pLEX_307-RhTRIM5-APEX2-V5; pDest40-HuTRIM5-APEX2-V5 and pLEX_307-HuTRIM5-APEX2-V5 were generated using Gateway recombination cloning. First, they were PCR amplified from available cDNA clones and recombined into pDONR221 using the BP reaction (Life Technologies, 11789-020) prior to being recombined into expression plasmids by LR cloning (Life Technologies, 11791-020). Plasmid constructs were verified by DNA sequencing. The AP1 luciferase reporter plasmid was a gift from Alexander Dent (Addgene plasmid #40342; 3XAP1pGL3), the NF-κB luciferase reporter was purchased from Promega (#E8491) and the Renilla luciferase plasmid (pRL-SV40, Addgene plasmid #27163) was a gift from Ron Prywes. Plasmid transfections were performed using Lipofectamine 2000 (ThermoFisher, 11668019) or Calcium Phosphate (Promega, E1200). Samples were prepared for analysis the day after DNA transfection.

### Antibodies and reagents

The following primary antibodies were used: V5 (Cell Signaling, 13202S), NCOA4 (Novus Biologicals, H00008031-M05), HA (Abcam, Ab9110), LRPPRC (Santa Cruz, 166178), FAF2 (Santa Cruz, 374098), NIPSNAP2 (Novus Biologicals, NBP2-45730), Prohibitin 2 (Novus Biologicals, NBP2-13754), Rad23B (Santa Cruz, 390019), TNIP1 (Cell Signaling, 4664), Calnexin (Santa Cruz, 23954), Calumenin (Santa Cruz, 271357), N4BP1 (Novus Biologicals, NBP2-37688), NIPSNAP1 (Santa Cruz, 515197), TBK1 (Cell Signaling, 3504S), Ube2D3 (Cell Signaling, 4330), Ube2N (Cell Signaling, 6999S), TIM23 (Santa Cruz, 514463), TOM20 (Santa Cruz, 17764), mCherry (Abcam, 183628), ATG13 (Cell Signaling, 13468S), GFP (Abcam, Ab290), FIP200 (Cell Signaling,12436 and Proteintech 17250-1-AP), MTCO2 (Abcam, 110258), VDAC1 (Abcam, 15895), DNA (Progen, AC-30-10), LC3B (Sigma, L7543; MBL International, PM036), TRIM5 (Cell Signaling #14326), Ubiquitin (MBL International, D058-3), Optineurin (Santa Cruz, 166576), NDP52 (Santa Cruz, 376540), ATG9 (Cell Signaling,13509) and actin (Santa Cruz, 58673). Secondary antibodies used were fluorescently conjugated goat anti-mouse (LI-COR, 925-68020) and goat anti-rabbit (LI-COR, 925-32210), HRP-conjugated goat anti-mouse (Bio-Rad, 1721011) and goat anti-rabbit (Bio-Rad, 1721019), Clean-Blot HRP (ThermoFisher, 21230), or Mouse TrueBlot ULTRA (Rockland, 18-8817).

Other reagents used were: nuclease free water (Dharmacon, B-003000-WB-100), Opti-MEM Reduced Serum Medium (ThermoFisher, 31985070), RIPA lysis buffer (ThermoFisher, 89901), phenylmethylsulfonyl fluoride (PMSF, Sigma, 93482, 1 mM), protease inhibitor cocktails (ROCHE, 11836170001), phospahatase inhibitor (ROCHE, 4906845001), BSA (Fisher Scientific, BP 1600-1), Puromycin (Sigma, P8833), Polybrene (EMD Millipore, TR-1003-G, 10 µg/ml), Hygromycin (Corning, 30-240-CR), MG132 (Selleckchem, S21619, 0.2 µM), IP lysis buffer (ThermoFisher, 87788), Dynabeads Protein G (ThermoFisher, 10004D), Streptavidin magnetic beads (ThermoFisher, 88816), and Restore PLUS Western blot stripping buffer (ThermoFisher, 46430). Trolox, CCCP, Oligomycin, Antimycin A, Ivermectin, and Propidium Iodide were purchased from Sigma.

### Western blotting, immunoprecipitation, and immunoflourescent labeling

SDS PAGE was carried out using pre-cast poly-acrylamide gels (Biorad) and immunoblotting was performed using standard procedures. Immunoprecipitation experiments were performed as previously described (Mandell et al., 2014). Immunoblot data was acquired using a Chemidoc MP instrument (Biorad) and quantitatively analyzed using Biorad Image Lab software. Immunoflourescent labeling of samples for high content imaging and confocal experiments, samples were fixed in 4% paraformaldehyde (Sigma) for 30 minutes prior to permeabilization in buffer containing 0.1% Saponin (Sigma) and 3% BSA. Following 1 hour incubation in primary antibodies, Alexafluor-conjugated secondary antibodies (Life Technologies) were used. Coverslips were mounted in ProLong Diamond anti-fade reagent (Life Technologies). Duolink *in situ* proximity ligation assays were performed according to the manufacturer’s instructions (Sigma).

### Luciferase assays

20000 HEK293T cells were plated in 96 well plates prior to transfection with the *Renilla* luciferase internal control reporter plasmid pRL-TK (thymidine kinase promoter dependent *Renilla* luciferase), plasmids encoding firefly luciferase responsive to NF-κB or AP1 APEX2 or TRIM5-APEX2 expression plasmids as previously described(Saha et al., 2020). 40-48h after transfection, the plate was assayed using the Dual-Glo Luciferase Assay System (Promega, E2920) and read using a Microplate Luminometer (BioTek, SYNERGY HTX Multi-Mode reader). Firefly luciferase readings were normalized to Renilla luciferase readings in each well, and the data are represented as fold-change compared to control cDNA.

### Infection assay using single-cycle viruses

8000 HeLa cells were seeded per well in a 96 well plate, 24h prior to virus challenge. Media containing VSV-G pseudotyped lentiviral vectors expressing GFP tagged HIV-1 (NL43, lacking Env) was added to infect cells in a total volume of 100 µL in the presence or absence of mitophagy inducers. After infection, cells were first incubated at 4°C for 1h to allow the virus to bind. Free virus was then removed by washing and cells were incubated in complete medium. 48h post infection, cells were fixed, stained with Hoechst 33342 and the fraction of transduced cells showing fluorescent protein positivity was determined by high content imaging and analysis as previously described(Saha et al., 2020).

### High content imaging

High content imaging and analysis were performed using a Cellomics CellInsight CX7 scanner equipped for live-cell imaging and driven by iDEV software (Thermo Fisher Scientific). Primary objects were cells (identified based on nuclear staining with DAPI or Hoechst 33342), and regions of interest (ROI) or targets were algorithm-defined by shape/segmentation, maximum/minimum average intensity, total area and total intensity to automatically identify puncta or other profiles within valid primary objects. Transduced cells were automatically identified based on having above background fluorescent protein signal in the nucleus. All data acquisition and analysis were computer driven and independent of human operators.

### APEX2-based biotin labeling and purification

Biotinylation reactions were performed as described previously (Jia et al., 2018). Briefly, HEK293T cells stably expressing APEX2-V5 or RhTRIM5-APEX2-V5 were cultured in the presence of 0.5 mM biotin-phenol (AdipoGen) for the last 30 minutes of the experiment. Cells were then pulsed with H2O2 (1 mM) for 1 minute prior to washing the cells with a buffer that quenches the biotinylation reactions (10 mM sodium ascorbate, 10 mM sodium azide, and 5 mM Trolox in PBS) and lysis in a buffer consisting of 6M urea, 0.3 M NaCl, 1 mM EDTA, 1 mM EGTA, 10 mM sodium ascorbate, 10 mM sodium azide, 5 mM Trolox, 1% glycerol and 25 mM TRIS-HCl (pH 7.5). Samples were then sonicated and insoluble material cleared by centrifugation. 3 mg of protein lysate per sample was then incubated with streptavidin-coated magnetic beads (Pierce) overnight. Beads were then washed with a series of buffers including: 1) IP lysis buffer (Pierce); 2) 1M KCl; 3) 50 mM Na2CO3; and 2M urea in 20 mM TRIS-HCl (pH 8.0).

### Sample preparation for LC-MS/MS

Protein samples on magnetic beads were washed four times with 200ul of 50mM Triethyl ammonium bicarbonate (TEAB) with a twenty-minute shake time at 4C in between each wash. Roughly 2.5 ug of trypsin was added to the bead and TEAB mixture and the samples were digested over night at 800 rpm shake speed. After overnight digestion the supernatant was removed and the beads were washed once with enough 50mM ammonium bicarbonate to cover. After 20 minutes at a gentle shake the wash is removed and combined with the initial supernatant. The peptide extracts are reduced in volume by vacuum centrifugation and a small portion of the extract is used for fluorometric peptide quantification (Thermo scientific Pierce). One microgram of sample based on the fluorometric peptide assay was loaded for each LC-MS analysis.

### Liquid Chromatography Tandem Mass Spectrometry

Peptides were desalted and trapped on a Thermo PepMap trap and separated on an Easy-spray 100 μm x 25 cm C18 column using a Dionex Ultimate 3000 nUPLC at 200 nL/min. Solvent A= 0.1% formic acid, Solvent B = 100% Acetonitrile 0.1% formic acid. Gradient conditions = 2% B to 50% B over 60 minutes, followed by a 50%-99% B in 6 minutes and then held for 3 minutes than 99% B to 2%B in 2 minutes and total run time of 90 minutes using Thermo Scientific Fusion Lumos mass spectrometer. The samples were run in DIA mode; mass spectra were acquired using a collision energy of 35, resolution of 30 K, maximum inject time of 54 ms and a AGC target of 50K, using staggered isolation windows of 12 Da in the m/z range 400-1000 m/z.

### Data Analysis

DIA data was analyzed using Spectronaut 15 (Biognosys Schlieren, Switzerland) using the directDIA workflow with the default settings. Briefly, protein sequences were downloaded from Uniprot (Human Proteome UP000005640), Human immunodeficiency virus type 1 (HIV-1) group M subtype B (isolate BRU/LAI; UP000007692), Tripartite motif-containing protein 5 from *Macaca mulatta* (Q0PF16), and common laboratory contaminant sequences from https://thegpm.org/crap/. Trypsin/P specific was set for the enzyme allowing two missed cleavages. Fixed Modifications were set for Carbamidomethyl, and variable modification were set to Acetyl (Protein N-term) and Oxidation. For DIA search identification, PSM and Protein Group FDR was set at 0.01%. A minimum of 2 peptides per protein group were required for quantification. A report was exported from Spectronaut using the reporting feature and imported into SimpliFi https://simplifi.protifi.com/ for QC and statistical analysis (Protifi, Farmingdale NY). Proteins ‘hits’ were defined as being enriched by 1.7-fold in the TRIM5-APEX2 samples (P < 0.05). Proteins known to be endogenously biotinylated were excluded from consideration. Gene set enrichment analysis of hits was performed using Reactome Pathway Browser (https://reactome.org/PathwayBrowser/#TOOL=AT). Mitochondrial proteins were identified using the MitoMiner4.0 database (https://mitominer.mrc-mbu.cam.ac.uk/release-4.0/begin.do), and gene ontology component analysis was performed using the Panther database (http://www.pantherdb.org/).

### Data availability

Mass Spectrometry Data is available at the MASSIVE data repository (massive.ucsd.edu) using number MSV000088006 and ProteomeExchange (http://www.proteomexchange.org) using ID number PXD028031.

### Mitochondrial isolation experiments

Subcellular fractionation was performed with a QProteome mitochondria isolation kit (Qiagen) according to the instruction manual. In brief, 10^7^ HeLa and Huh7 cells were re-suspended in 1 mL of lysis buffer, incubated for 10 min at 4⁰ C and centrifuged at 1000x g for 10 min. The supernatant was transferred into a separate tube as cytosolic fraction, while the pellet was re-suspended in 1.5 mL of ice-cold disruption buffer, rapidly passed through 26g needle 10-15 times to disrupt cells and centrifuged at 1000x g for 10 min, 4⁰ C. The supernatant was then re-centrifuged at 6000x g for 10 min, 4⁰ C. The pellet obtained after centrifugation comprised the mitochondrial fraction. For proteinase K (PK) digestion, mitochondria were re-suspended in Mitochondrial buffer (MB) (210 mM mannitol, 70 mM sucrose, 10 mM HEPES, 1 mM EDTA, pH 7.5) with 50 mg/mL of PK and incubated 30 min at RT. For Triton X-100 digestion, mitochondria were re-suspended in digestion buffer (10 mM sucrose, 0.1 mM EGTA/Tris and 10 mM Tris/HCl, pH 7.4) with Triton X-100. Both reactions were stopped by addition of 5 mM phenylmethylsulfonyl fluoride (Sigma). For the analysis of integral membrane proteins, the mitochondrial fraction was re-suspended in MB buffer or MB buffer containing freshly prepared 0.1 M Na2CO3 (pH 11.5) and incubated on ice for 30 min. The insoluble membrane fraction was centrifuged at 16000x g for 15 min.

### Confocal microscopy

Sub-airy unit (0.6AU) pinhole confocal microscopy with a Zeiss LSM800 or a Leica TCS-SP8 microscope was performed followed by computational image restoration with Huygens Essential (Scientific Volume Imaging, Hilversum, Netherlands) utilizing a constrained maximum likelihood estimation algorithm. All 3D images were acquired with a 63X/1.4NA plan apochromat oil immersion objective and sampled at ideal Nyquist sampling rates in x, y, and z planes. Voxel lateral and axial dimensions were determined by utilizing an online Nyquist calculator (https://svi.nl/NyquistCalculator) allowing for sub-diffraction limited resolution following image restoration. All images were rendered on a high performance CUDA-GPA enabled workstation and 3D renders were generated for volume-object analysis with Huygens Object Analyzer and Colocalization software.

### Metabolic profiling

We used the Seahorse XF Cell Mito Stress Test Kit (Agilent) according to the manufacturer’s protocol in a Seahorse XFe96 Analyzer to measure oxygen consumption and extracellular acidification rates in Huh7 cells treated or not with 1.5 µM IVM. After initial optimization experiments, we used kit components at the following concentrations: oligomycin, 1.5 µm; FCCP, 1 µm; rotenone, 0.5 µm; antimycin A, 0.5 µm. Analysis was performed using Seahorse Analytics software (Agilent).

### Cell death analysis

WT and TRIM5 knockout Huh7 cells were treated or not with 20 µM IVM for 24 hours prior to measurement of cell viability by measuring propidium iodide (PI) labeling or using the alamarBlue cell viability assay (Invitrogen). For PI experiments, cells were harvested and treated with 1 µg/ml PI and 10 µg/ml Hoechst 33342 prior to flow cytometry using an Attune NxT flow cytometer (Thermo). PI staining intensity was measured in single cells as determined by forward and side scatter and by Hoechst positivity. 20k total cells were analyzed per treatment. For the alamarBlue assays, reagent was added to cells 20 h prior to determining absorbance at 570 nm and 600 nm using a SYNERGY HTX Multi-Mode Reader (BioTek) according to the manufacturer’s protocol.

### Statistical analysis

Data are expressed as means ± SEM (n>3). Data were analyzed with unpaired two-tailed t-tests or ANOVA with Bonferroni post hoc analysis. Statistical significance is defined as *, P < 0.05; **, P < 0.01; ***, P < 0.001; ****, P < 0.0001.

## SUPPLEMENTARY FIGURE LEGENDS

**Figure S1.**
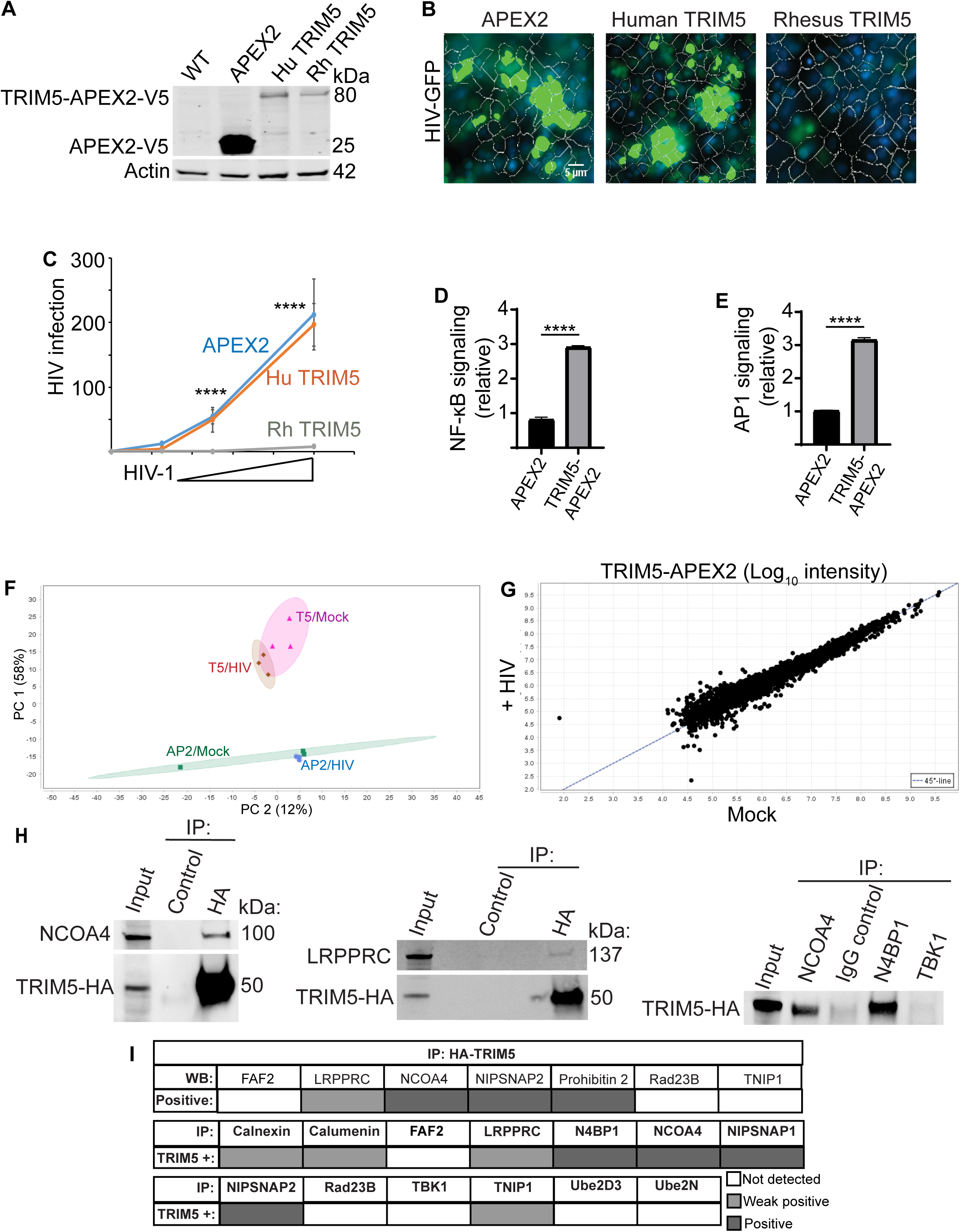
Identification of TRIM5-proximal proteins by mass spectrometry and immunoprecipitation. **(A)** Immunoblot analysis of APEX2-V5 or TRIM5-APEX2-V5 expression in stably transduced HEK293T cells. **(B,C)** High content imaging based analysis of HIV restriction in APEX2-V5 expressing cell lines. The indicated stably transduced cell line was exposed to different dilutions of VSV-G pseudotyped HIV-1 encoding a GFP reporter. 2 days post infection, cells were fixed and stained with Hoechst 33342. Cells were automatically identified by nuclear staining and the above-background GFP intensity was measured per cell. N = 6 replicates with > 500 cells per replicate analyzed. Data, mean ± SEM; ****, *P* < 0.0001 by ANOVA. **(D, E)** Impact of TRIM5-APEX2 expression on NF-κB and AP1 signaling in HEK293T cells transfected with plasmids encoding firefly luciferase under the control of NF-κB- or AP1- responsive promoters, constitutively active *Renilla* luciferase, and either TRIM5-APEX2 or APEX2 alone. 40-48h after transfection, samples were harvested, and luciferase values determined. Data, mean + SEM; ****, *P* < 0.0001 by Student’s t test. **(F)** Principle component analysis proteomic data from cells stably expressing rhesus TRIM5-APEX2-V5 (T5) or APEX2-V5 alone (AP2) and infected with VSV-G pseudotyped HIV-1 (HIV) or not (mock) for 3 hours prior to sample preparation. Shaded regions show 95% confidence ellipses. **(G)** Scatter plot of proteomic data from TRIM5-APEX2-V5 expressing cells infected or not with VSV-G. Dashed 45⁰ line indicates no difference between samples. **(H)** Coimmunoprecipitation analysis of interactions between TRIM5 and selected proteins identified as TRIM5 proximal in mass spectrometry data from HeLa cells stably expressing HA-tagged TRIM5. **(I)** Summary of coimmunoprecipitation results from data shown in this figure and throughout the manuscript. All proteins tested were identified as being enriched in TRIM5-APEX2 datasets. Lysates from HeLa cells stably expressing TRIM5-HA were subjected to immunoprecipitation with anti-HA and immunoblots probed with the indicated antibodies (top matrix) or immunoprecipitated with the antibodies indicated in the middle and bottom matrices and immunoblots probed with anti-HA.

**Figure S2.**
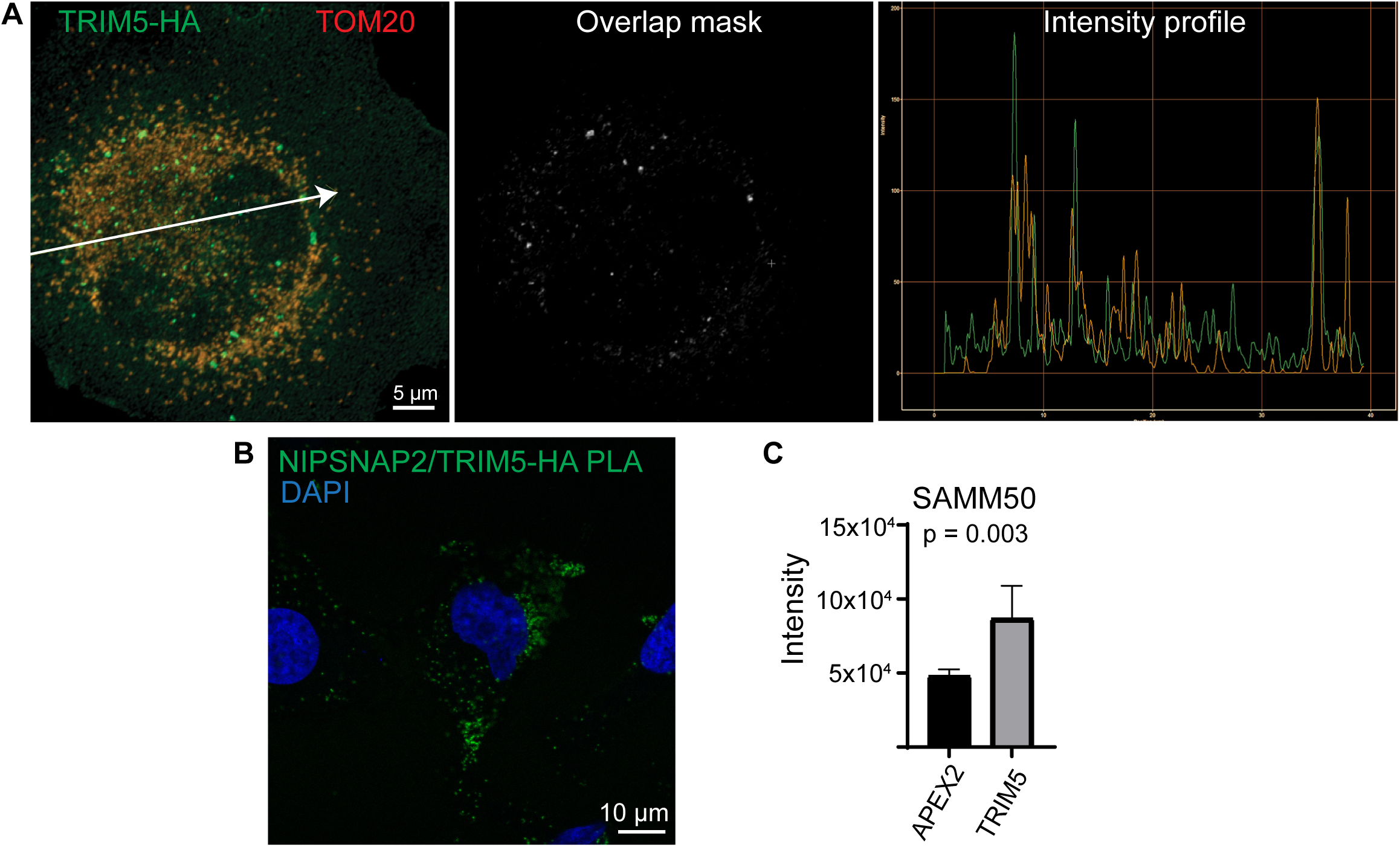
TRIM5 can localize to mitochondria and interact with the mitophagy receptor NIPSNAP2. **(A)** Confocal microscopic analysis of colocalization between mitochondrial marker TOM20 and TRIM5-HA in HeLa cells. Left, maximum image projection of a representative deconvolved confocal image. Arrow illustrates a transect through the cell along which the fluorescence intensity was measured in the TRIM5-HA (green) and TOM20 (red) channels as shown on the right. The middle image shows regions in which TRIM5 and TOM20 signals overlap in X, Y, and Z planes. **(B)** Proximity ligation assay (PLA) demonstrating that anti- NIPSNAP2 and anti-HA antibodies recognize antigens within 16 nm of each other in HeLa cells stably expressing TRIM5-HA. Positive PLA signal results in the formation of green fluorescent puncta. **(C)** SAMM50 intensity measurements from mass spectrometry based quantitation of biotinylated proteins from APEX2 and RhTRIM5-APEX2 cells. Data, mean + SEM. *P* value determined by t test.

**Figure S3.**
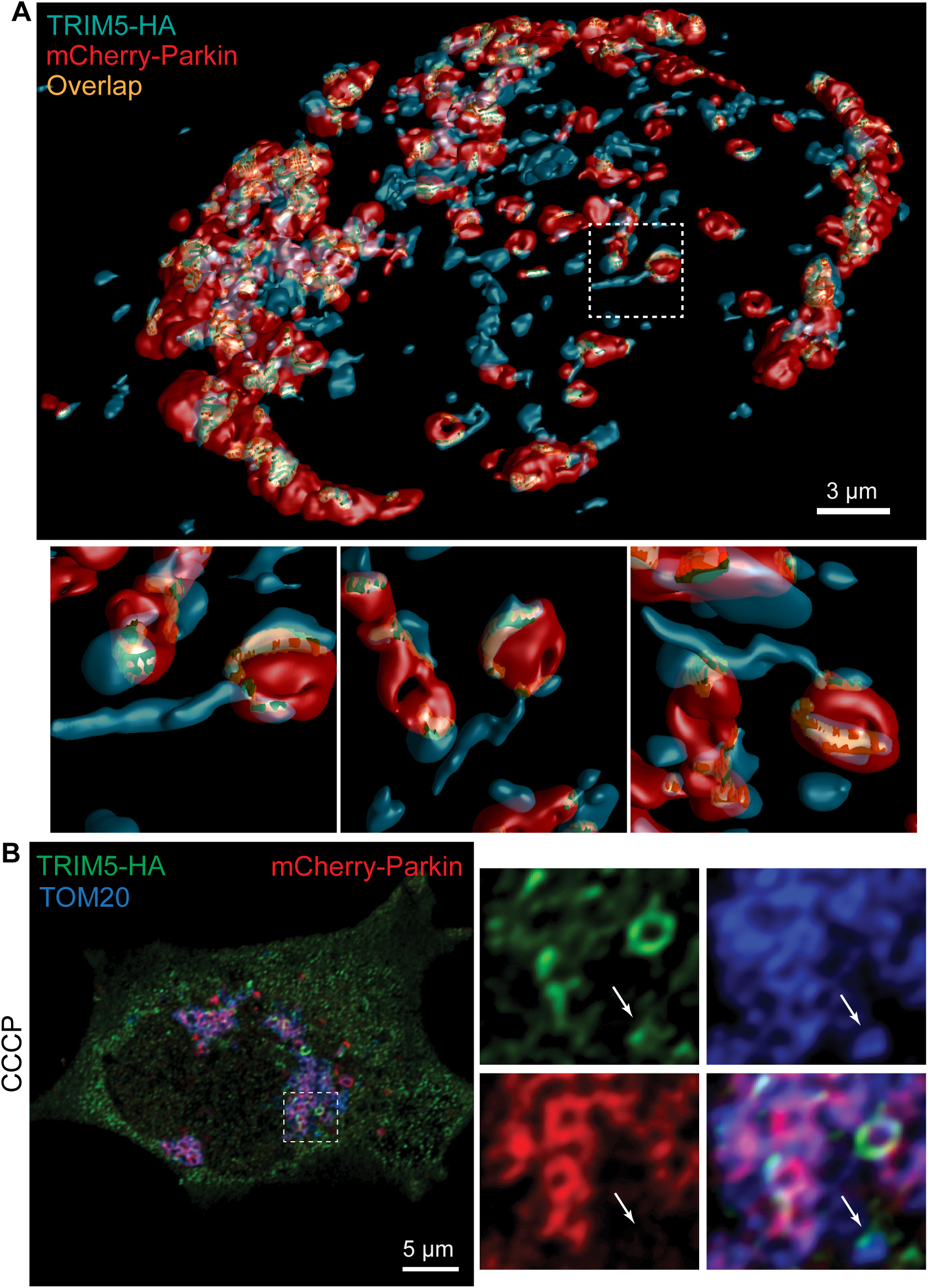
Localization of TRIM5 in cells following mitochondrial uncoupling with CCCP. HeLa cells stably expressing TRIM5-HA were transiently transfected with mCherry-tagged Parkin and treated with 20 µM CCCP for 3 h prior to fixation. **(A)** 3D reconstruction of cells stained with antibodies recognizing HA. Zoomed in images of the region demarcated by the dashed line box are shown below. Each image shows the same structures from different viewpoint in X, Y, and Z planes. Voxels showing overlapping TRIM5 and Parkin signal are indicated by the gold color. **(B)** Deconvolved confocal image of a CCCP-treated cell stained to detect TRIM5-HA, TOM20, and mCherry-Parkin. Zoomed in images of the boxed region are shown to the right. Arrow indicates a TRIM5-HA positive mitochondrion that is negative for mCherry-Parkin.

**Figure S4.**
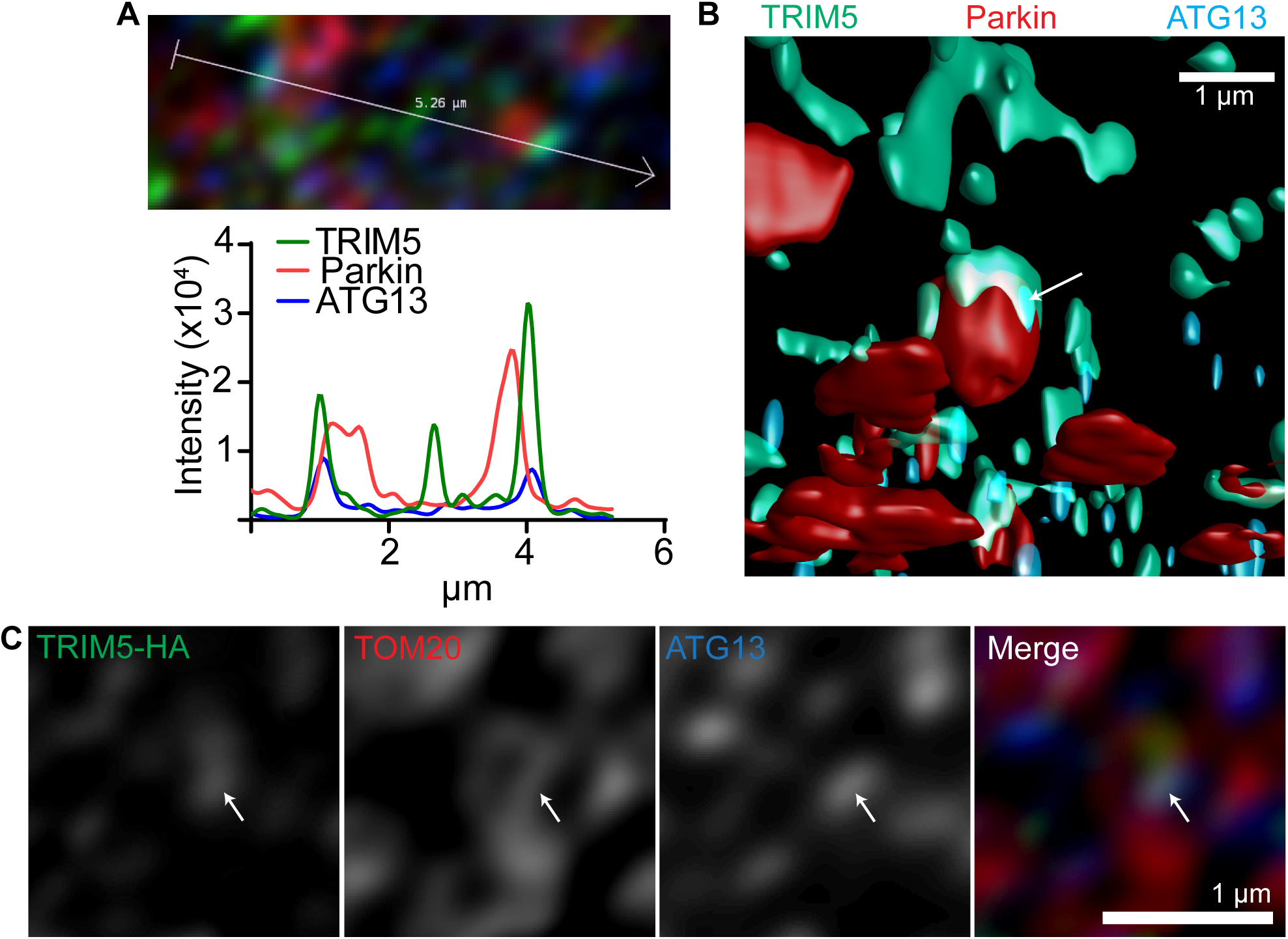
ATG13 and TRIM5 colocalize on the surface of damaged mitochondria. **(A)** Intensity profile from a deconvolved confocal image of a cell stained to detect TRIM5-HA (green), mCherry-Parkin (red), and endogenous ATG13 (blue) showing colocalized TRIM5 and ATG13 signals at specific foci on mCherry-Parkin positive mitochondria following a 3 hour treatment with CCCP. **(B)** 3D reconstruction of a cell stained as in (A). Arrow indicates a TRIM5-ATG13 double-positive region on the outer face of a Parkin-positive mitochondrion. **(C)** Localization of TRIM5 to ATG13-positive foci on mitochondria does not require Parkin. HeLa cells stably expressing TRIM5-HA (but not transfected with Parkin) were treated with CCCP for 3 hours prior to fixation and staining with antibodies recognizing HA, TOM20, and ATG13. Image shows a deconvolved confocal image. Arrow indicates colocalized TRIM5 and ATG13 signal on the outside of a mitochondrion.

**Figure S5.**
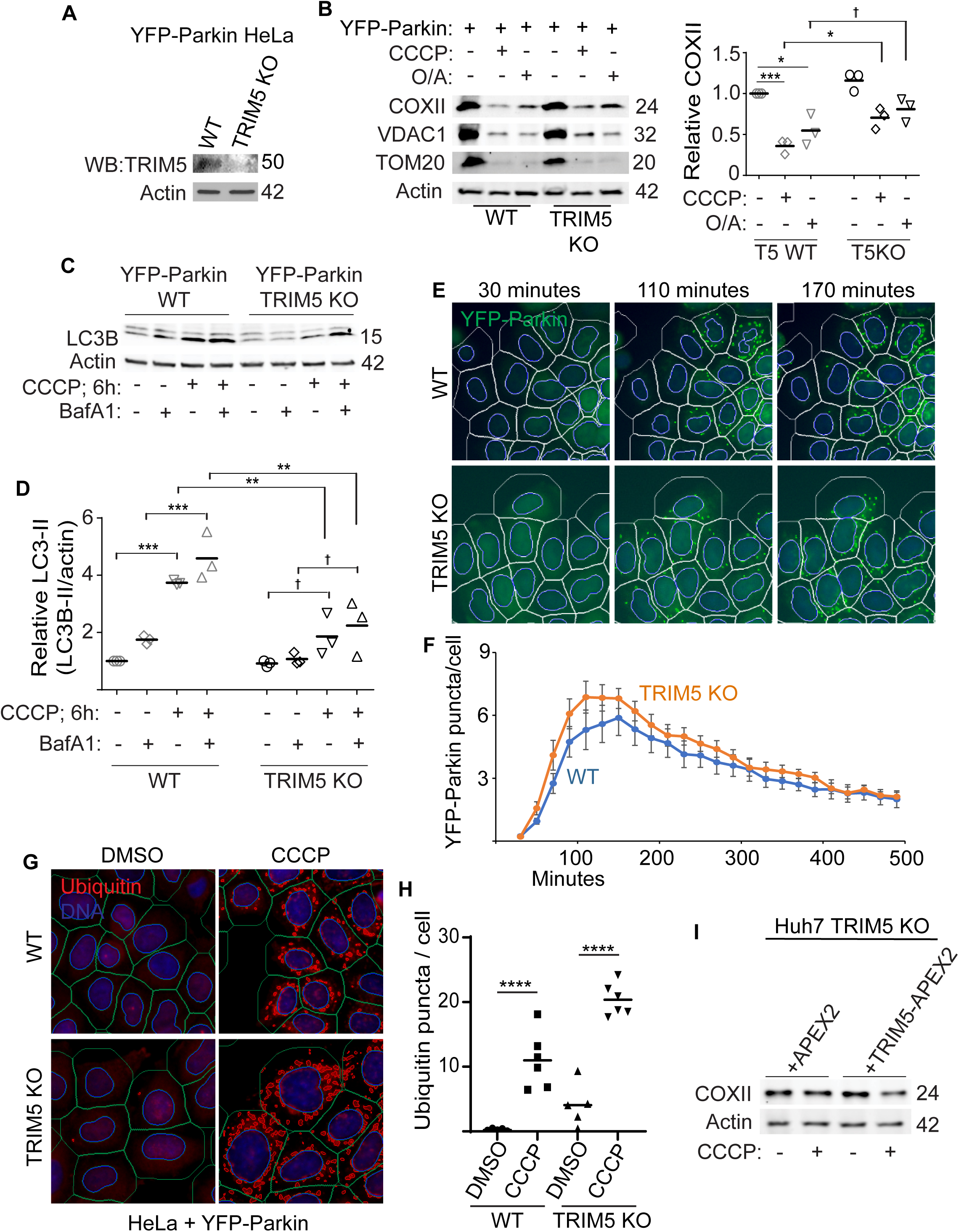
Impacts of TRIM5 knockout in a HeLa cell model of Parkin-dependent mitophagy. **(A)** Immunoblot analysis of TRIM5 expression in HeLa cells stably expressing YFP-Parkin following transduction with lentivirus expressing Cas9 and either non-targeting or TRIM5-targeting guide RNA. **(B)** Immunoblot analysis of the effect of CCCP or oligomycin/antimycin (O/A) treatment on the abundance of mitochondrial proteins in WT and TRIM5 knockout HeLa cells that stably express YFP-Parkin. Data: *, *P* < 0.05; ***, *P* < 0.001; †, not significant by ANOVA. N = 3 independent experiments. **(C,D)** Immunoblot analysis of the impact of TRIM5 knockout on CCCP-induced autophagy in YFP-Parkin expressing HeLa cells treated or not with Bafilomycin A1 (BafA1). Data: **, *P* < 0.01; ***, *P* < 0.001; †, not significant by ANOVA. N = 3 independent experiments. **(E, F)** Time-lapse high content imaging analysis of YFP-Parkin puncta abundance in WT and TRIM5 knockout cells following treatment with 20 µM CCCP for the indicated time points. Blue ring and white rings in E, automatically generated nuclear and cell boundary masks. Yellow mask, machine-identified YFP-Parkin puncta. Data, mean ± SEM. N = 6 independent experiments with >500 cells analyzed per experiment. **(G,H)** High content imaging and analysis of the abundance of ubiquitin puncta in WT and TRIM5 KO HeLa cells expressing YFP-Parkin and treated or not with 20 µM CCCP for 3 hours. G, representative images of cells stained to detect poly-ubiquitin (red) and DNA (blue). Blue and green rings show automatically masked nuclear and cellular boundaries. Red mask indicates coalesced ubiquitin bodies identified within the cytoplasm. Data: ****, *P* < 0.0001 by ANOVA. N = 5 experiments with >500 cells analyzed per experiment. **(I)** The impact of CCCP treatment on the abundance of the mitochondrial protein COXII in TRIM5 knockout Huh7 cells in which APEX2-tagged TRIM5 had been reintroduced by lentiviral transduction. Transduction with an APEX2-alone expressing lentivirus was used as a negative control.

**Figure S6.**
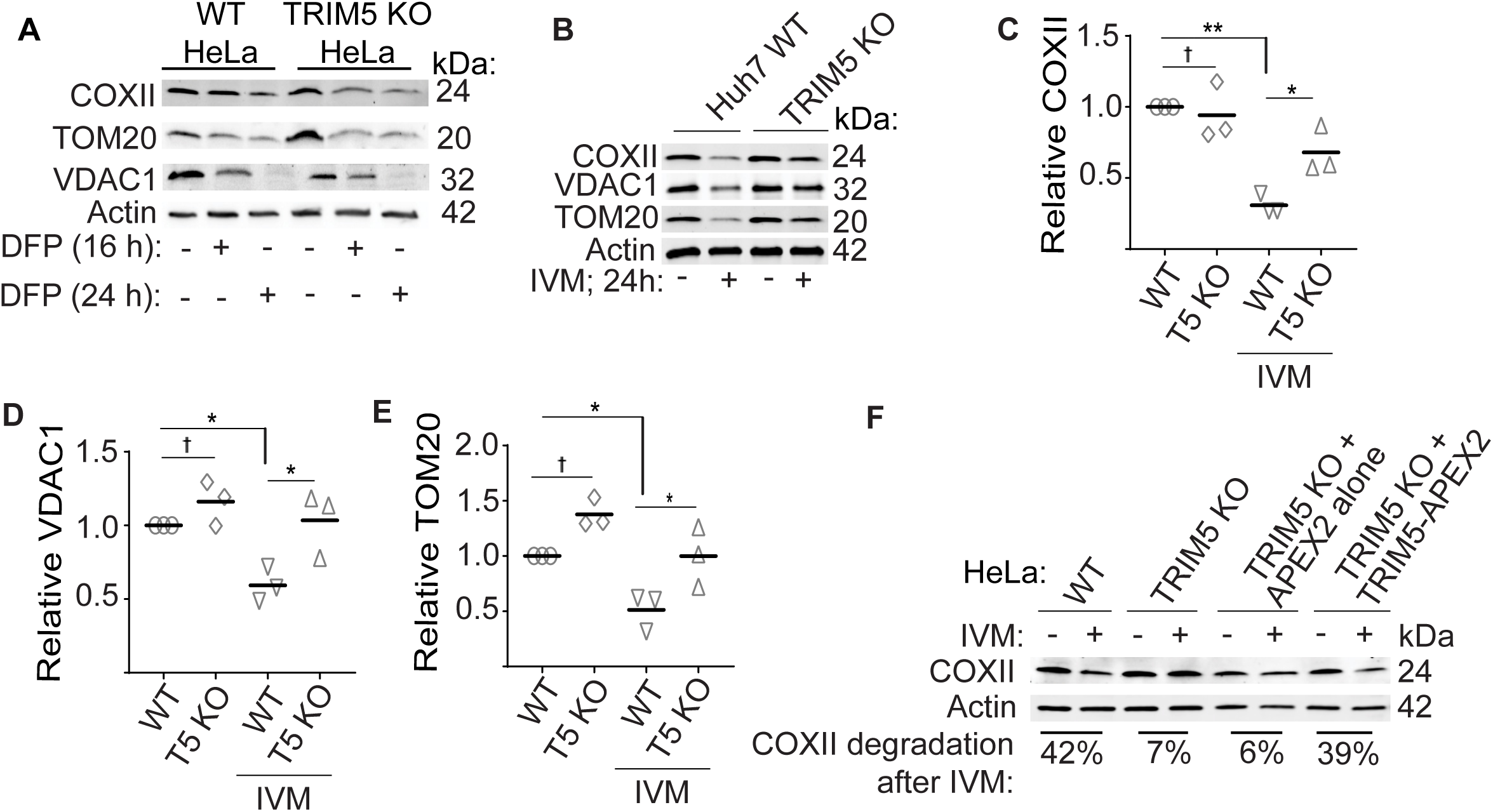
Role of TRIM5 in Parkin-independent mitophagy. **(A)** Immunoblot analysis of the impact of DFP on the abundance of the indicated mitochondrial proteins in whole cell lysates from WT and TRIM5 knockout HeLa cells. Actin is used as a loading control. **(B-E)** Immunoblot analysis of the effect of ivermectin (IVM) treatment on the abundance of mitochondrial proteins in WT and TRIM5 knockout Huh7 cells. Cells were treated with IVM for 24 hours prior to lysis and immunoblotting with the indicated antibodies. Plots, the abundance of mitochondrial proteins COXII, VDAC1, and TOM20 was determined relative to actin. Data: *, *P* < 0.05; **, P < 0.01; †, not significant by ANOVA. N = 3 independent experiments. **(F)** The impact of IVM treatment on the abundance of COXII in WT and TRIM5 knockout HeLa cells stably expressing TRIM5-APEX2 or APEX2 alone. Percentages represent the amount of COXII (relative to actin loading control) lost after IVM treatment for each cell line.

**Figure S7.**
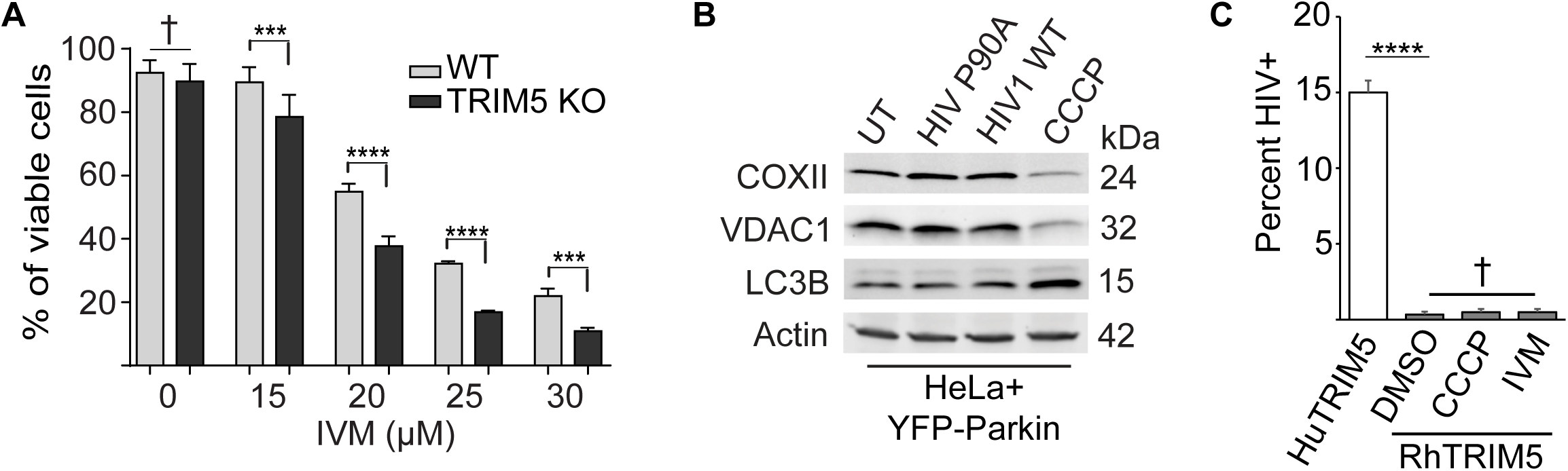
TRIM5-dependent mitophagy is independent of TRIM5 actions in HIV restriction. **(A)** AlamarBlue-based assessment of cell viability in WT and TRIM5 knockout Huh7 cells following 24 h treatment with the indicated concentrations of IVM. **(B)** The impact of infection with TRIM5-sensitive (HIV P90A) or TRIM5-resistant (HIV WT) pseudoviruses on mitochondrial protein abundance and on LC3B conversion in HeLa cells expressing YFP-Parkin. Cells were subjected to synchronous infection with VSV-G pseudotyped HIV for 24 hours prior to lysis and immunoblotting with the indicated antibodies. CCCP treatment (24 hours) was used as a positive control for mitophagy induction. **(C)** The impact of mitophagy-inducing compounds on the ability of RhTRIM5 to restrict HIV-1. HeLa cells stably expressing HuTRIM5 or RhTRIM5 were infected with VSV-G pseudotyped HIV-1 encoding a GFP reporter after 1.5 hours pre-treatment with CCCP, IVM, or DMSO control. Virus was then allowed to enter cells for 4 h in the presence of the compounds, after which the media was replaced. 48 h later, the percent of cells showing GFP positivity was determined by high content imaging. Data, mean ± SEM; ***, *P* < 0.001; ****, *P* < 0.0001; †, not significant by ANOVA.

